# Cysteamine-bicalutamide combination treatment restores alpha-ketoglutarate and corrects proximal tubule phenotype in cystinosis

**DOI:** 10.1101/2020.02.10.941799

**Authors:** Amer Jamalpoor, Charlotte AGH van Gelder, Fjodor A Yousef Yengej, Esther A Zaal, Sante Princiero Berlingerio, Koenraad R Veys, Carla Pou Casellas, Koen Voskuil, Khaled Essa, Carola ME Ammerlaan, Laura Rita Rega, Reini van der Welle, Marc R Lilien, Maarten B Rookmaaker, Hans Clevers, Judith Klumperman, Elena Levtchenko, Celia R Berkers, Marianne C Verhaar, Maarten Altelaar, Rosalinde Masereeuw, Manoe J Janssen

## Abstract

Nephropathic cystinosis is a severe monogenetic kidney disorder caused by mutations in *CTNS*, encoding the lysosomal transporter cystinosin, resulting in lysosomal cystine accumulation. The sole treatment, cysteamine, slows down the disease progression, but does not correct the established proximal tubulopathy. Here, we developed a new therapeutic strategy by applying an omics-based strategy to expand our knowledge on the complexity of the disease and prioritize drug targets in cystinosis. We identified alpha-ketoglutarate as a key metabolite linking cystinosin loss, lysosomal autophagy defect and proximal tubular impairment in cystinosis. This insight offered a bicalutamide-cysteamine combination treatment as a novel dual target pharmacological approach for the phenotypical correction of cystinotic proximal tubule cells, patient-derived kidney tubuloids and cystinotic zebrafish.

## Introduction

Nephropathic cystinosis (MIM219800) is a lysosomal storage disease (LSD) caused by mutations in *CTNS*, a gene that codes for the lysosomal cystine/proton symporter cystinosin ^1^. The loss of cystinosin leads to the lysosomal accumulation of cystine throughout the body and causes irreversible damage to various organs, particularly the kidneys ^2^. The first clinical signs develop during infancy (age of ~6 months) in the form of renal Fanconi syndrome, a generalized proximal tubular dysfunction leading to chronic kidney failure ^3^. For the past decades, great efforts have been directed towards reducing cellular cystine accumulation in cystinotic patients. The cystine-depleting drug cysteamine can efficiently lower the lysosomal cystine levels and postpone disease progression. However, it poses serious side effects and does not correct the established proximal tubulopathy associated with cystinosis ^3,4^.

Recent discoveries in lysosomal gene expressions and their regulation networks has provided new insights in the studies of LSDs, including cystinosis ^5–7^. Several studies based on *in vitro* and *in vivo* models have demonstrated that the loss of cystinosin is indeed associated with disrupted lysosomal autophagy dynamics ^8^, accumulation of distorted mitochondria ^9^ and increased reactive oxygen species (ROS) ^10^, leading to abnormal proliferation and dysfunction of proximal tubule cells. Regardless of the observed cellular defects associated with cystinosis, the mechanism linking cystinosin loss, lysosomal defects and epithelial dysfunction remains unknown, hampering the development of an enduring intervention to combat the disease.

In this work, we aimed to develop a novel pharmacological strategy to treat cystinosis by correcting its renal tubular phenotype. By combining CRISPR/Cas9 technology and a dual proteomics and metabolomics approach, we identified multiple pathway disruptions in *CTNS* deficient renal proximal tubule cells. Particularly, we identified alpha-ketoglutarate (αKG), an important intermediate of the tricarboxylic acid (TCA) cycle, as part of a unifying mechanism linking cystinosin loss, lysosomal autophagy disruption and proximal tubular impairment in cystinosis. Using this knowledge, we identified the bicalutamide-cysteamine combination treatment as a novel strategy for the phenotypical correction of cystinotic proximal tubule cells. This approach was further validated in cystinotic patient-derived kidney tubuloids and cystinotic zebrafish, demonstrating a therapeutic potential for this combination therapy to treat patients with cystinosis.

## Results

### CRISPR-generated *CTNS^−/−^* ciPTEC display increased cystine accumulation and impaired lysosomal autophagy dynamics

A guide RNA (gRNA) targeting exon 4 of the *CTNS* gene was used to introduce mutations by CRISPR/Cas9 in conditionally immortalized proximal tubule cells (ciPTEC). After cell sorting and subsequent clonal cell expansion, three clones with biallelic mutations (line 3, 7, and 35) were initially selected and all displayed a similar phenotype (Supplementary fig. S1a-d), therefore, line 3 (hereafter referred to as *CTNS^−/−^*) was used for subsequent experiments. This model enables direct evaluation of the effect of *CTNS* loss on proximal tubular cells independent of chronic exposure to other disease related changes in the body. As a reference, we also included a non-isogenic patient-derived *CTNS^−/−^* ciPTEC line bearing the homozygous 57-kb deletion ^11,12^, which will be referred to as *CTNS^Patient^*. In accordance with the pathophysiology of the disease, *CTNS^−/−^* cells displayed significantly increased levels of cystine compared to control *CTNS^WT^* cells (5.19 ± 0.30 vs. 0.05 ± 0.02 nmol/mg protein), comparable to *CTNS^Patient^* cells (Fig. 1a).

**Fig. 1.**
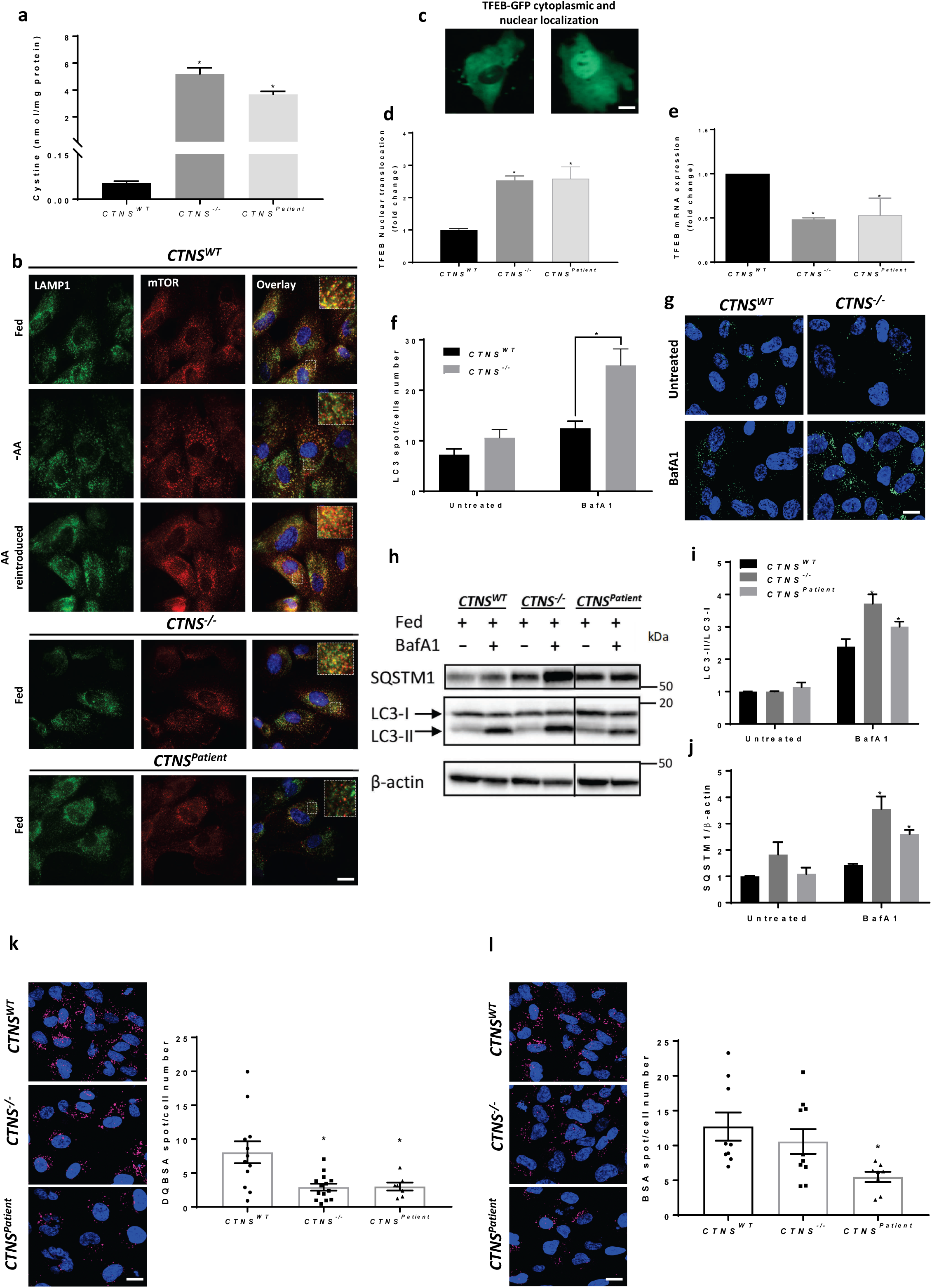
CRISPR-generated *CTNS^−/−^* ciPTEC demonstrate lysosomal cystine accumulation and distorted lysosomal autophagy dynamics. **a**, Quantification of cystine levels (nmol/mg protein) by HPLC-MS/MS in control (*CTNS^WT^*), CRISPR-generated cystinotic cells (*CTNS^−/−^*), and patient-derived cystinotic cells (*CTNS^Patient^*). **b**, Representative immunofluorescent staining of *CTNS^WT^*, *CTNS^−/−^*, and *CTNS^Patient^* co-immunolabelled with lysosomal-associated membrane protein 1 (LAMP1; green) and mTOR (Red). Merge images with zoomed areas are representative of the localization of mTOR with lysosomes in various experimental conditions. Scale bars are 13 µm**. c**, **d**, Representative confocal micrographs and quantification of TFEB-GFP nuclear translocation in *CTNS^WT^*, *CTNS^−/−^*, and *CTNS^Patient^*, respectively. Scale bars are 20 µm. **e**, *TFEB* mRNA expression of the *CTNS^−/−^* and *CTNS^Patient^* cells was measured using real-time quantitative polymerase chain reaction, with ribosomal protein S13 (RPS-13) as a reference gene, normalized to control cells. **f**, **g**, Quantification of LC3-II accumulation and representative confocal micrographs in *CTNS^WT^* and *CTNS^−/−^* cells in presence and absence of 25 nM bafilomycin (BafA1) for 4 hrs, respectively. Scale bars are 20 µm. **h**, **i**, **j**, Western blotting and densitometric analyses for LC3-II/LC3-I ratio and SQSTM1 protein levels in *CTNS^WT^*, *CTNS^−/−^*, and *CTNS^Patient^* cells cultured in the presence or in the absence of 25 nM BafA1 for 4 h, respectively. β-Actin was used as a loading control. **k**, **l**, Representative confocal micrographs and quantification of DQ-BSA and BSA in *CTNS^WT^*, *CTNS^−/−^*, and *CTNS^Patient^* cells, respectively. Scale bars are 20 µm. Data are expressed as the mean ± SEM of three independent experiments. Statistical analysis was performed using one way analysis of variance (ANOVA) followed by Dunnett’s multiple comparisons test. P-values < 0.05 were considered to be significant.

Next we evaluated the effect of *CTNS* loss on mammalian target of rapamycin complex 1 (mTORC1)-mediated autophagy ^8,9,13^. Under normal conditions, mTORC1, is bound to the lysosomes and is responsible for regulating a wide range of cellular processes, including autophagy ^14,15^. In the presence of nutrients (standard medium, fed condition), mTOR was located on the lysosomal membranes of *CTNS^WT^* cells (Fig. 1b). Upon starvation (-AA) mTOR was released from the lysosomes and re-localized upon reintroduction of nutrients. In contrast, in *CTNS^−/−^* and *CTNS^Patient^* cells mTOR was already released from lysosomes in the fed condition (Fig. 1b), indicating dissociation of the mTOR complex in cystinosis. We further evaluated mTOR activity in the cells by tracking the subcellular localization of transcription factor EB (TFEB). If mTOR is deactivated, unphosphorylated TFEB can translocate to the nucleus (Fig. 1c), where it regulates gene transcription and activates autophagy. A ~2.5-fold increase in TFEB nuclear translocation was observed after transfection with TFEB-GFP in *CTNS^−/−^* and *CTNS^Patient^* cells compared to *CTNS^WT^* cells (Fig. 1d). As TFEB will downregulate its own expression after activation ^16^, we found that the endogenous *TFEB* mRNA expression was also reduced in *CTNS^−/−^* and *CTNS^Patient^* cells (2-fold), when compared to control cells (Fig. 1e). During autophagy LC3-II is recruited to autophagosomes and p62/SQSTM1 is degraded after fusion of autophagosomes with the lysosomes ^17^. Blocking the lysosomal fusion with bafilomycin showed a significant increase in both LC3-II and p62/SQSTM1 levels in *CTNS^−/−^* and *CTNS^Patient^* cells compared to *CTNS^WT^* cells (Fig. 1f-j), indicating an abnormal induction of autophagy with delayed lysosomal cargo degradation in cystinotic cells. This was further confirmed when *CTNS^−/−^* and *CTNS^Patient^* cells showed a reduced ability (~2.5-fold) to process bovine serum albumin conjugated with a BODIPY dye (DQ BSA), a dye which is endocytosed by the cells and become fluorescent after degradation inside the lysosomes, compared to control cells (Fig. 1k). *CTNS^Patient^* cells, but not *CTNS^−/−^* cells, demonstrated a decreased BSA uptake compared to the control cells (Fig. 1l).

### Metabolomic and proteomic profiling reveal alpha-ketoglutarate accumulation in cystinosis

To gain in-depth knowledge of the affected cellular pathways and proximal tubule impairments in cystinosis, we performed targeted metabolomic profiling of 100 key metabolites (Supplementary file 1) and untargeted proteomics in the three cell lines (Fig. 2). Principal component analysis (PCA) of the metabolites and over 4500 identified proteins (Fig. 2a, f) showed that *CTNS^Patient^* cells account for most of the variability in the data, indicating that the different genetic background of the *CTNS^Patient^* cells affect the data more than the *CTNS* loss itself. This was further visualized by unsupervised hierarchical clustering in which the isogenic *CTNS^−/−^* cells cluster with *CTNS^WT^* rather than *CTNS^Patient^* cells (Fig. 2b, supplementary fig. S2a). To explore which pathways are directly linked to *CTNS* loss we therefore focused on the metabolites and proteins that were significantly altered in *CTNS^−/−^* cells compared to the *CTNS^WT^*. Pathway enrichment analysis of the metabolites distinctively affected in *CTNS^−/−^* cells revealed numbers of affected pathways, including cystine/cysteine metabolism, the TCA cycle, glutathione (GSH) metabolism, and amino acid metabolisms (Fig. 2c). Proteomic profiling, on the other hand, revealed a total of 337 proteins that were up- or downregulated in *CTNS^−/−^* cells compared to the control cells (Fig. 2g). The differentially abundant proteins were then subjected to gene ontology (GO) classification via the Panther Classification System database ^18^ to highlight their molecular role in the cell (Fig. 2h). The analysis showed an overall reduction in proteins involved in lipid metabolism and the breakdown of micromolecular cellular structures (peptidases, amino acid catabolism, reductases and hydrolases), and an increase in the glycolysis, TCA cycle, DNA replication and DNA repair, in line with the metabolomics.

**Fig. 2.**
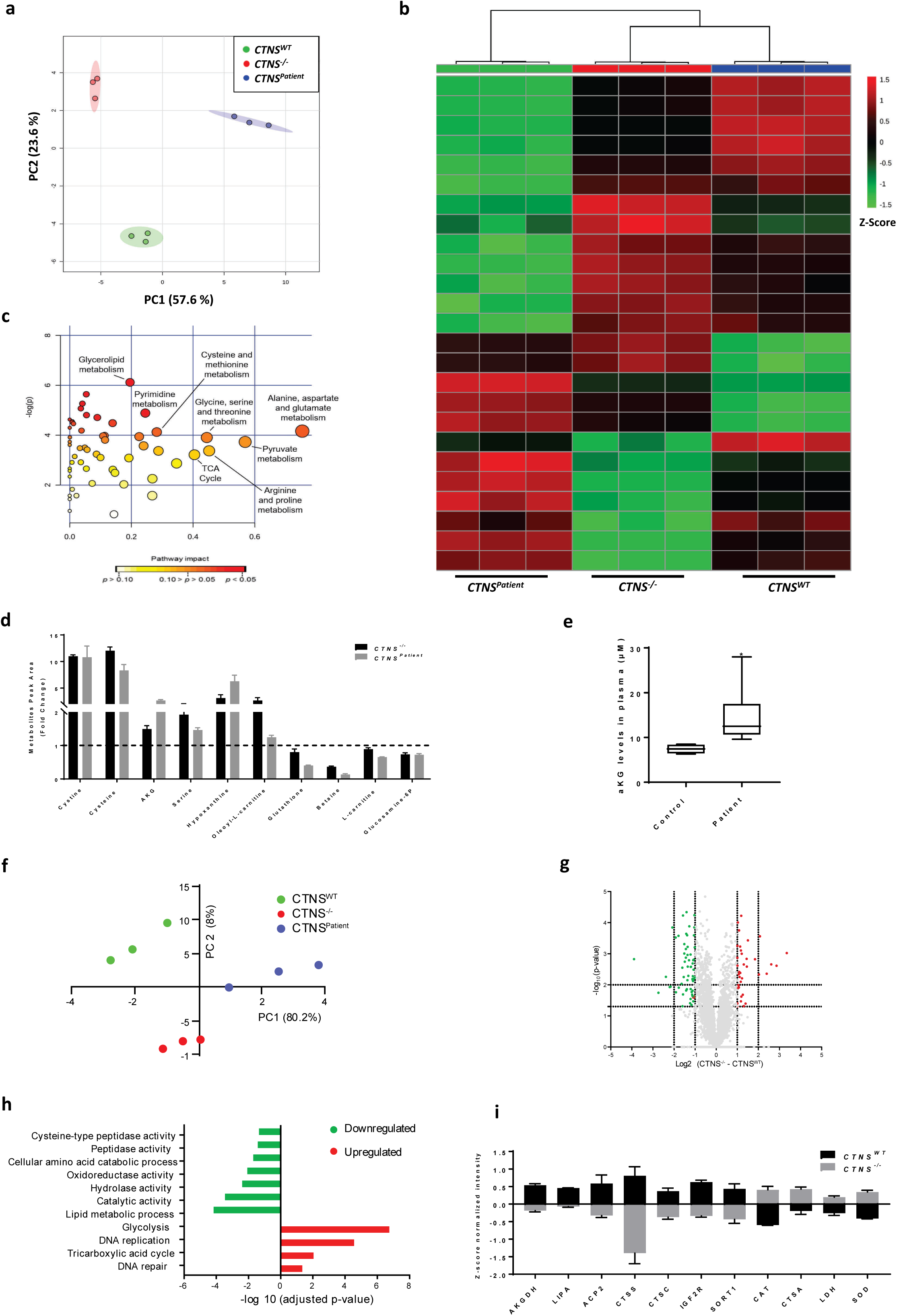
Metabolomic and proteomic profiling reveal alpha-ketoglutarate accumulation in cystinosis. **a**, Principal component analysis (PCA) of control (*CTNS^WT^*), CRISPR-generated cystinotic cells (*CTNS^−/−^*), and patient-derived cystinotic cells (*CTNS^Patient^*) based on the metabolites measured. Each dot represents one sample, and the dots of the same color are biological replicates. **b**, Heatmap analysis of top 25 metabolites distinctively expressed in control and *CTNS^−/−^* proximal tubule cells. The rows display different metabolites and the columns represent the samples. The Z-score of each feature is plotted in red-green color scale. Metabolites significantly decreased (P < 0.05) were displayed in green, while metabolites significantly increased (P < 0.05) were displayed in red. **c**, Global test pathway enrichment analysis of the intracellular metabolic interactions distinctively affected in *CTNS^−/−^* cells compared to healthy control cells. Larger circles further from the y-axis and orange-red color show higher impact of pathway affected in *CTNS^−/−^* cells. **d,** List of metabolites that were shared and significantly upregulated or downregulated in both the *CTNS^−/−^* and *CTNS^Patient^* cells compared to control cells. Data are presented as the fold change of three independent experiments performed in triplicate. **e,** Plasma levels of αKG (µM) in healthy individuals (n=4) and cystinotic patients (n=6). All cystinotic patients were on cysteamine treatment at the time of blood sampling. **f,** Principal component analysis (PCA) of the measured proteins in *CTNS^WT^*, *CTNS^−/−^*, and *CTNS^Patient^*. **g,** Volcano plot illustrates significantly differentially abundant proteins. The -log_10_ (Benjamini–Hochberg corrected P-value) is plotted against the log_2_ (fold change: *CTNS^WT^*/*CTNS^−/−^*). The non-axial vertical lines denote ±1.5-fold change while the non-axial horizontal line denotes P=0.05, which is our significance threshold (prior to logarithmic transformation). **h,** Gene ontology (GO) analysis illustrates classes of proteins differing between *CTNS^WT^* and *CTNS^−/−^* cells. Proteins with significant differences between the two cell types were subjected to GO classification via the Panther Classification System database in terms of molecular function. **i,** List of proteins that were significantly upregulated and downregulated in *CTNS^−/−^* cells compared to control cells. LIPA; Lysosomal acid lipase, ACP2; Lysosomal acid phosphatase, AKGDH; Alpha-ketoglutarate dehydrogenase, CTSS; Cathepsin S, CTSC; Dipeptidyl peptidase, IGFR2; Mannose-6-phosphate receptor, SORT1; Sortilin, CAT; Catalase, CTSA; Lysosomal protective protein, and SOD; Superoxide dismutase. Data are expressed as the mean ± SEM. Statistical analysis was performed using unpaired two-tailed Student’s t-test. P-values < 0.05 were considered to be significant.

When cross-checking the affected metabolites from both *CTNS^−/−^* and *CTNS^Patient^* cells, we identified a set of key metabolites that were significantly changed as a result of *CTNS* loss: cystine, cysteine, αKG, serine, hypoxanthine, and oleoyl-L-carnitine levels were increased, whereas the levels of GSH, betaine, L-carnitine, and glucosamine-6-phosphate were decreased in both *CTNS^−/−^* and *CTNS^Patient^* compared to control cells (Fig. 2d). αKG was of particular interest, as αKG is known to play a central role in the TCA cycle, maintaining the redox balance, regulating mTOR and autophagy. Moreover, αKG is a well-known antioxidant and its level can be increased by cells in response to oxidative stress contributing to the prevention and/or treatment of several disorders induced by oxidative stress ^19–21^. This is done through downregulation of the mitochondrial enzyme AKGDH, reported to be severely diminished in human pathologies where oxidative stress is thought to play a vital role ^21^. Indeed we found a decreased expression of AKGDH in *CTNS^−/−^* cells (Fig. 2i). We also observed a significantly increased level of αKG in plasma of cystinotic patients compared to healthy controls (Fig. 2e; Table 1), signifying that loss of *CTNS* also leads to an increase in aKG levels in patients. This prompted us to further evaluate the effect of increased levels of αKG in cystinosis pathology.

**Table 1.**
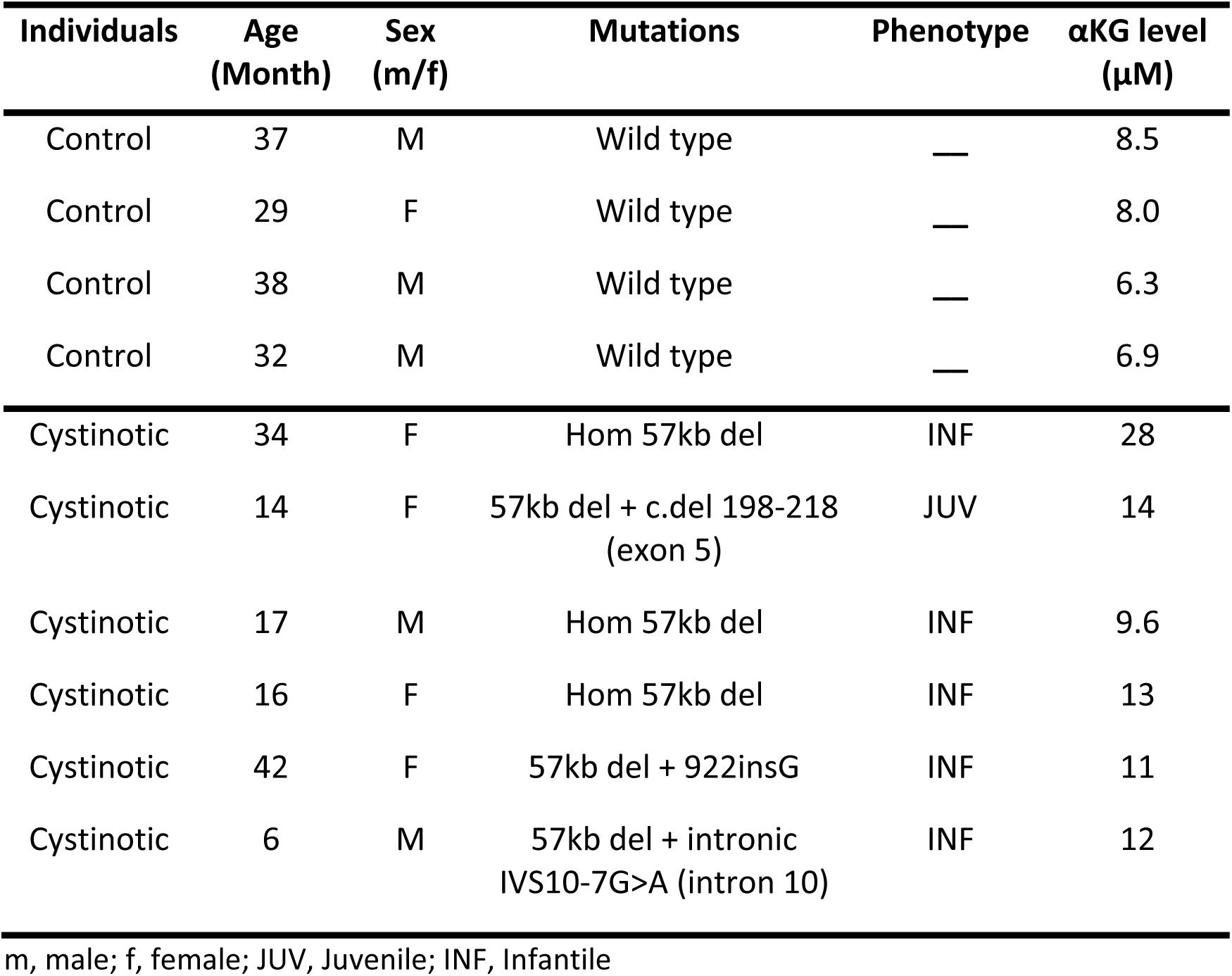
Alpha ketoglutarate levels in plasma of control and cystinotic individuals.

### Alpha-ketoglutarate regulates the phenotypic alterations in cystinosis

The link between cystinosin loss-of-function and increased oxidative stress have been long-established ^10,11,22,23^. In line with increased αKG levels in response to the oxidative stress, both *CTNS*^−/−^ and *CTNS^Patient^* cells presented with increased ROS levels (~1.5-fold), when compared to healthy control cells (Fig. 3a). Starvation further triggered ROS production in all the three cell lines (Fig. 3b-d). Supplementing starved cells with dimethyl αKG (DMKG), a cell permeable form of αKG for 4 hrs led to a reduction in ROS levels in *CTNS^WT^* cells (2.1-fold), a modest reduction in *CTNS*^−/−^ cells (1.2-fold), but increased ROS levels in *CTNS*^Patient^ cells (Fig. 3b-d), showing the anti-oxidative effect of αKG in *CTNS^WT^* but not in *CTNS* deficient cells. Cystinotic cells were also more sensitive to starvation as seen by an increase in cell death and caspase positive cells after 24 hrs (Fig. 3e-g). Adding DMKG during the 24 hrs starvation had little to no effect on control cells, but led to massive cell death and apoptosis activation in cystinotic cells (Fig. 3e-g, and supplementary fig. S3a-c). In line with previous studies ^24–26^, we therefore hypothesized that the increased level of αKG could be the main mechanism perturbing autophagy and therefore promoting cell death in cystinotic proximal tubule cells. DMKG exposure indeed resulted in an increased level of LC3-II/LC3-I ratio in *CTNS^−/−^* and *CTNS^Patient^* but not in control cells, confirming that αKG is a key metabolite abnormally inducing autophagy in cystinotic cells (Fig. 3h, i). In light of our presented data, we identified αKG as a key metabolite linking cystinosin loss, increased oxidative stress, autophagy disruption, and proximal tubule dysfunction in cystinosis (Fig. 3j).

**Fig. 3.**
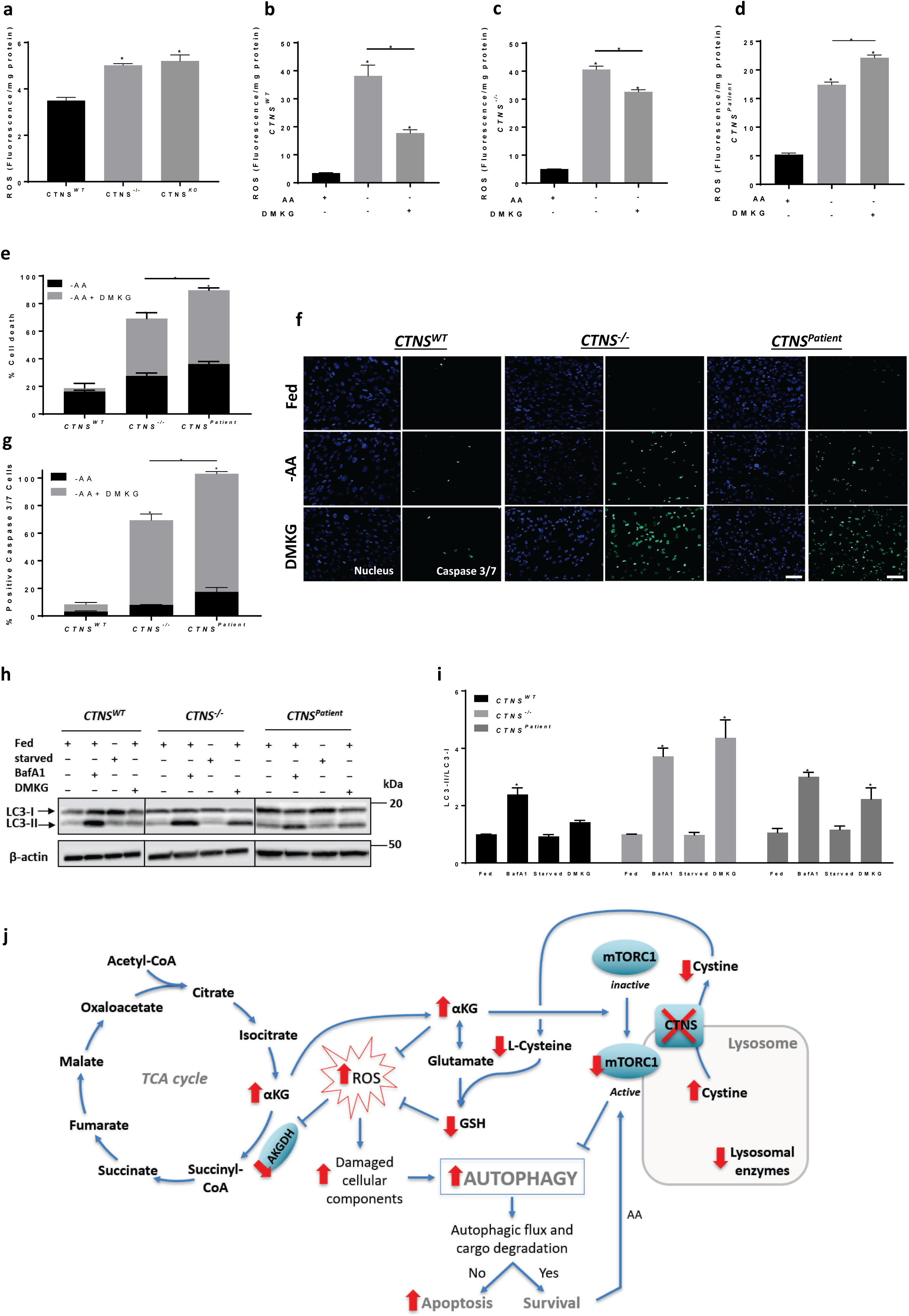
Alpha-ketoglutarate is a key metabolite responsible for impaired autophagy and proximal tubule dysfunction in *CTNS* deficient renal proximal tubule cells. **a,** Relative reactive oxygen species (ROS/mg protein) production in *CTNS^WT^*, *CTNS^−/−^*, and *CTNS^Patient^* cells. **b**, **c**, **d**, Relative reactive oxygen species (ROS/mg protein) production in *CTNS^WT^*, *CTNS^−/−^*, and *CTNS^Patient^* cells upon starvation for 4 hrs in the presence and absence of DMKG (4 hrs), respectively. **e**, Cell viability test of DMKG (2 mM)-treated control (*CTNS^WT^*), CRISPR-generated cystinotic cells (*CTNS^−/−^*), and patient-derived cystinotic cells (*CTNS^Patient^*) in fed and starved condition (-AA) for 24 hrs. **f, g,** Representative confocal micrographs (Scale bars are 20 µm) and immunofluorescence analysis of caspase 3/7 activation in DMKG (2 mM)-treated *CTNS^WT^*, *CTNS^−/−^*, and *CTNS^Patient^* cells in fed and starved condition for 24 hrs, respectively. **h**, **i**, Western blotting and densitometric analyses for LC3-II/LC3-I ratio in *CTNS^WT^*, *CTNS^−/−^*, and *CTNS^Patient^* cells cultured in the presence or in the absence of BafA1 (25 nM) and DMKG (2 mM) for 4 hrs, respectively. β-Actin was used as a loading control. **j**, Working model summarizing the results obtained in this work. TCA cycle; tricarboxylic acid cycle, αKG; alpha-ketoglutarate, AKGDH; alpha-ketoglutarate dehydrogenase, ROS; Relative reactive oxygen species, GSH; glutathione, mTORC1; mammalian target of rapamycin complex 1, AA; Amino acid. Data are expressed as the mean ± SEM of three independent experiments. Statistical analysis was performed using one way analysis of variance (ANOVA) followed by Dunnett’s multiple comparisons test. Where appropriate unpaired two-tailed Student’s t-test were used. P-values < 0.05 were considered to be significant.

### Bicalutamide and cysteamine combination treatment phenotypically corrects *CTNS^−/−^* proximal tubule cells

Cysteamine is the only drug used to efficiently reduce cystine accumulation in patients with cystinosis. However, it cannot correct the established proximal tubulopathy associated with the disease. We hypothesized that this may be linked to the inability of this drug to lower αKG and target other metabolic pathways associated with *CTNS* loss. We therefore screened different candidate drugs based on their ability to reduce cystine and αKG levels and restore the metabolic profile using metabolomics (Supplementary file 1). The concentrations of the drugs tested were within a non-cytotoxic range (Supplementary fig. S3d-j). Cysteamine, as expected, boosted GSH and lowered cystine and cysteine levels. However, it had no significant impact on the other metabolites or on the proteome profiles (Fig. 4a-e, supplementary fig. S4a). Among the candidate drugs tested, bicalutamide, an anti-androgenic agent, did not restore the high cystine and cysteine levels but could improve the overall metabolic phenotype, including αKG (Fig. 4a). Bicalutamide alone also could modestly upregulate several metabolic enzymes involved in cysteine conversion, lysosomal degradation, and the TCA cycle including AKGDH enzyme (Fig. 4c-e, supplementary fig. S4b). The cysteamine-bicalutamide combination treatment, however, showed an additive effect and unsupervised clustering shows that the cells receiving the combination therapy are more similar to the *CTNS^WT^* cells than the other conditions (Fig. 4b). Furthermore, combination treatment of *CTNS^−/−^* cells led to the upregulation of more than 50 proteins, revealing an enrichment of proteins involved in the TCA cycle, and in the metabolism of macromolecules such as lipids and vitamins that were shown to be initially downregulated in *CTNS^−/−^* cells (Fig. 4c-e, supplementary fig. S4c).

**Fig. 4.**
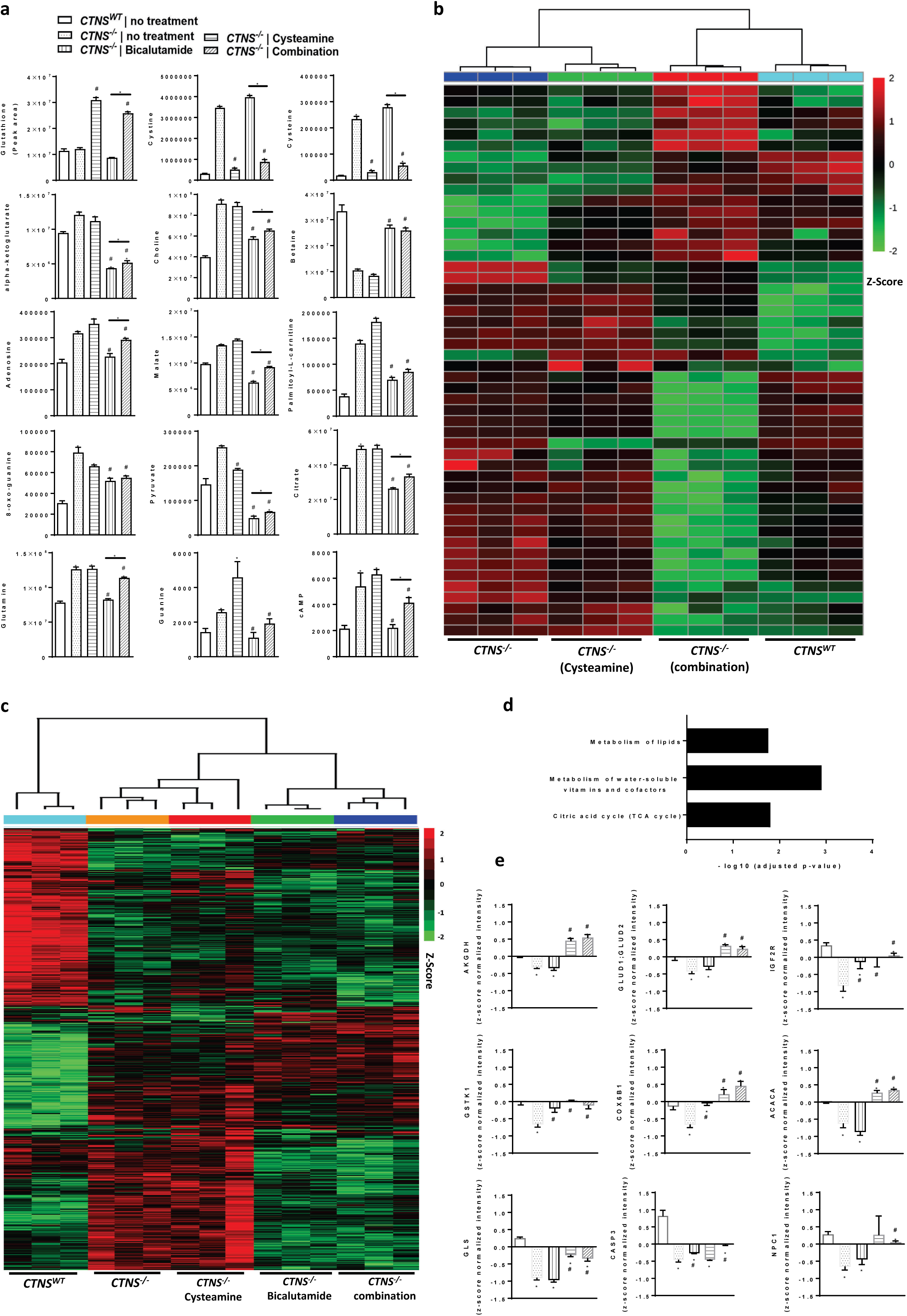
Bicalutamide-cysteamine combination treatment: A novel treatment correcting the metabolome and proteome profile of *CTNS* deficient renal proximal tubule cells. **a**, Metabolomic analysis of CRISPR-generated cystinotic cells (*CTNS^−/−^*) treated with cysteamine (100 µM), bicalutamide (35 µM), and a combination of cysteamine and bicalutamide (100 µM and 35 µM, respectively). **b**, Heatmap analysis of the measured metabolites in *CTNS^−/−^* cells upon treatment with cysteamine and cysteamine-bicalutamide combination treatment. The row displays metabolite feature and the column represents the samples. The row Z-score of each feature is plotted in red-green color scale. Metabolites significantly decreased were displayed in green, while metabolites significantly increased were displayed in red. **c**, **d**, Heatmap and REACTOME analysis of the altered proteins in *CTNS^−/−^* cells upon treatment with cysteamine, bicalutamide, and cysteamine-bicalutamide combination treatment. The row displays protein feature and the column represents the samples. The row Z-score of each feature is plotted in red-green color scale. Proteins significantly decreased were displayed in green, while metabolites significantly increased were displayed in red. **e**, Proteomic analysis of *CTNS^−/−^* cells treated with cysteamine (100 µM), bicalutamide (35 µM), and a combination of cysteamine and bicalutamide (100 µM and 35 µM, respectively), respectively. AKGDH; Alpha-ketoglutarate dehydrogenase, GLUD1; GLUD2; Mitochondrial glutamate dehydrogenase 1/2, IGF2R; Mannose-6-phosphate receptor, GSTK1; Glutathione S-transferase kappa 1, COX6B1; Cytochrome c oxidase subunit 6B1, ACACA; Acetyl-CoA carboxylase 1-Biotin carboxylase, GLS; Glutaminase kidney isoform, mitochondrial, CASP3; Caspase-3, NPC1; Niemann-Pick C1 protein. Data are expressed as the mean ± SEM. Statistical analysis was performed using one way analysis of variance (ANOVA) followed by Dunnett’s multiple comparisons test. Where appropriate unpaired two-tailed Student’s t-test were used. * Significantly different from *CTNS^WT^* cells (p < 0.05). # significantly different from *CTNS^−/−^* cells (p < 0.05).

As bicalutamide reduced the elevated level of αKG in cystinotic cells, we hypothesized that it could also resolve αKG-mediated downstream effects. Indeed, bicalutamide (but not cysteamine) reduced the αKG-mediated cell death and αKG-mediated increase in LC3-II/LC3-I ratio in *CTNS^−/−^* cells (Fig. 5a, b). Although cysteamine alone showed no effect on autophagy, combined with bicalutamide additively reduced the αKG-mediated increase in LC3-II/I in *CTNS*^−/−^ cells compared to bicalutamide alone (Fig. 5b). Notably, cysteamine, bicalutamide, and their combination had no effect on the basal autophagy activity (Supplementary fig. S4d).

**Fig. 5.**
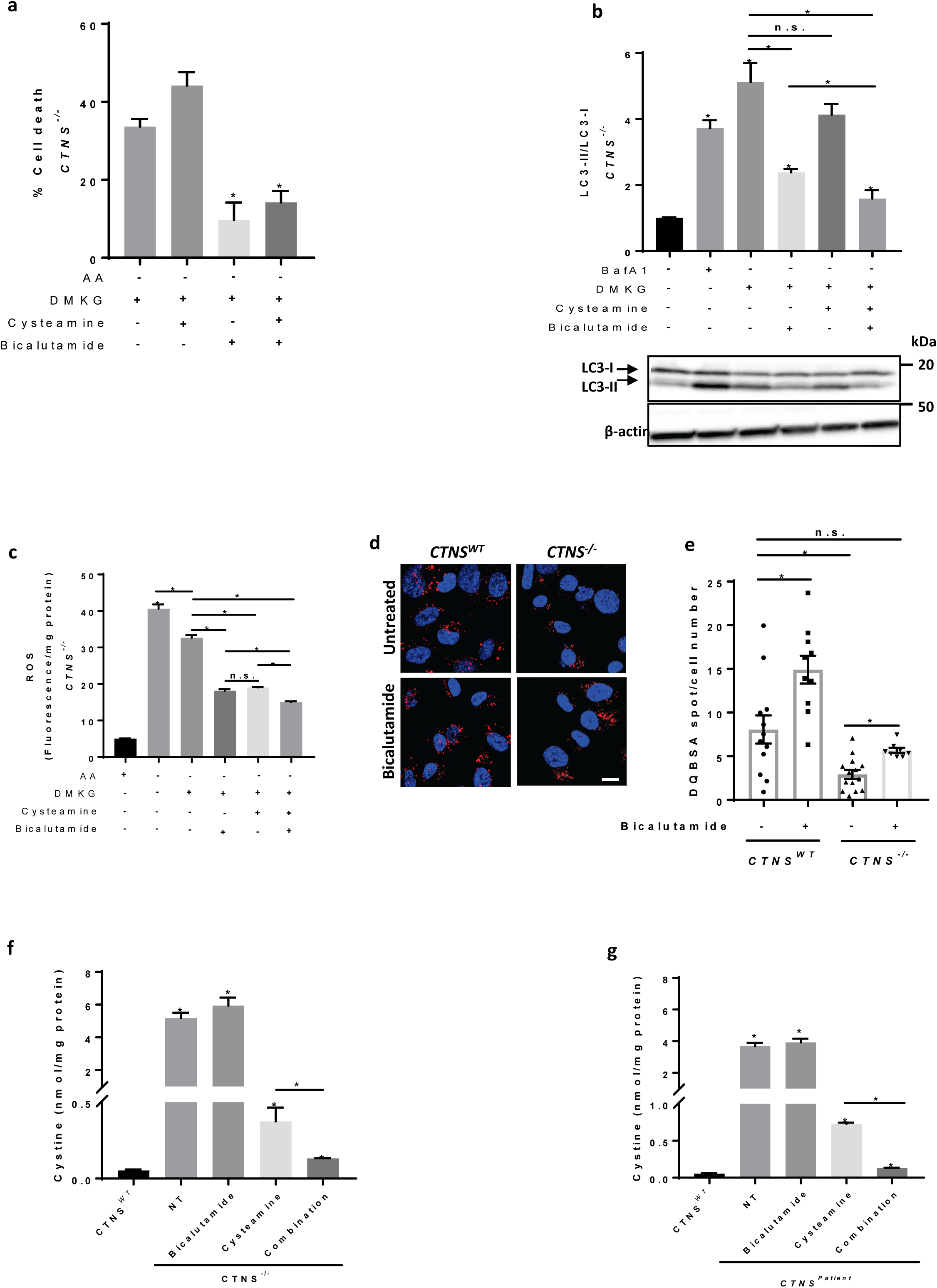
Cysteamine-bicalutamide combination treatment efficiently lowers lysosomal cystine, abolishes αKG-mediated autophagy distortion and cell death in *CTNS* deficient renal proximal tubule cells. **a**, Cell viability test of DMKG (2 mM)-treated CRISPR-generated cystinotic cells (*CTNS^−/−^*) upon pretreatment with cysteamine (100 µM), bicalutamide (35 µM), and a combination of cysteamine and bicalutamide (100 µM and 35 µM, respectively). **b**, Western blotting and densitometric analyses for LC3-II/LC3-I ratio in DMKG (2 mM)-treated *CTNS^WT^* and *CTNS*^−/−^ cells upon pretreatment with cysteamine (100 µM), bicalutamide (35 µM), and a combination of cysteamine and bicalutamide (100 µM and 35 µM, respectively). β-Actin was used as a loading control. **c**, Relative reactive oxygen species (ROS/mg protein) production in DMKG-treated *CTNS^WT^* and *CTNS^−/−^* cells upon pretreatment with cysteamine (100 µM), bicalutamide (35 µM), and a combination of cysteamine and bicalutamide (100 µM and 35 µM, respectively). **d**, **e**, Representative confocal micrographs and quantification of DQ-BSA in *CTNS^WT^*, and *CTNS^−/−^* cells upon treatment with bicalutamide (35 µM). Scale bars are 20 µm. **f**, **g**, Quantification of cystine levels (nmol/mg protein) by HPLC-MS/MS in *CTNS^−/−^*, *CTNS^Patient^* in the absence of the drug (NT) or upon treatment with cysteamine (100 µM), bicalutamide (35 µM), and a combination of cysteamine and bicalutamide (100 µM and 35 µM, respectively), respectively. Data are expressed as the mean ± SEM of three independent experiments. Statistical analysis was performed using one way analysis of variance (ANOVA) followed by Dunnett’s multiple comparisons test. Where appropriate unpaired two-tailed Student’s t-test were used. P-values < 0.05 were considered to be significant.

We next tested whether bicalutamide alone or in combination with cysteamine could retrieve the antioxidant effect of αKG in cystinotic cells. Similar to cysteamine, bicalutamide was able to retrieve the effect of αKG in *CTNS*^−/−^ cells (2.1-fold; Fig. 5c). Further reduction in ROS levels was observed when bicalutamide and cysteamine were given together. Of note, bicalutamide alone had no effect on GSH levels (Fig. 4a) but could significantly lower the ROS levels in presence of αKG in cystinotic cells (Fig. 5c), suggesting that bicalutamide, similar to cysteamine through a yet unknown mechanism, retrieves the antioxidant property of αKG in *CTNS^−/−^* cells.

Recently, bicalutamide has been patented for the treatment of several LSDs, promoting mTOR-associated autophagy and TFEB-mediated cellular exocytosis ^27^, and therefore allowing the relief and/or treatment of the symptoms of many LSDs. Accordingly, bicalutamide was able to induce TFEB nucleus translocation in both control and *CTNS*^−/−^ cells and downregulate *TFEB* mRNA expression in control cells (Supplementary fig. S4e, f). In agreement with the increased TFEB activity, bicalutamide also restored endocytic cargo processing in cystinotic cells (Fig. 5d, e), allowing us to test whether the increased degradation of lysosomal cargo results in the decreased cystine accumulation. We used a more sensitive and quantitative LC-MS/MS method to measured cystine levels in cystinotic cells upon treatment with cysteamine, bicalutamide, or their combination. In line with metabolomics data, *CTNS*^−/−^ cells treated with bicalutamide alone did not exhibit any reduction in cystine levels. However, cysteamine alone resulted in a reduction of cystine in *CTNS*^−/−^ cells (0.37 ± 0.08 nmol/mg protein; Fig. 5f). However, this was still higher (7.5- fold) than that found in control cells (0.05 ± 0.02 nmol/mg protein). Interestingly, combination treatment of bicalutamide and cysteamine resulted in a 2.8-fold decrease in cystine when compared to cysteamine alone (0.13 ± 0.003 *vs* 0.37 ± 0.08 nmol/mg protein; Fig. 5f), bringing lysosomal cystine close to the control. This effect was also found in *CTNS^Patient^* cells (Fig. 5g).

### Bicalutamide and cysteamine combination treatment efficiently lowers cystine and αKG levels in patient-derived cystinotic kidney tubuloids

Next, we evaluated the safety and efficacy of cysteamine-bicalutamide combination treatment in patient-derived tubuloids, an advanced *in vitro* model of cystinosis ^28^. We established kidney tubuloids from the urine of two pediatric cystinotic patients (*CTNS^Patient-1^* and *CTNS^Patient-2^*) and compared their characteristics to tubuloids derived from healthy kidney tissue from two donors (*CTNS^WT-1^ and CTNS^WT-2^*). Urine-derived tubuloids were positive for paired-box gene 8 (PAX8) and negative for tumor protein p63 (Fig. 6a), confirming that these structures indeed consist of kidney epithelium (PAX8+/p63-) and not urothelium (PAX8-/p63+). Moreover, both patient and control tubuloids robustly expressed markers of various segments of the nephron consisted of proximal tubule, loop of Henle, distal tubule and collecting duct (Supplementary fig. S5a), recapitulating the optimal physiological situation ^28^. Both *CTNS^Patient^* tubuloids displayed cystinosis phenotype of increased cystine accumulation (~7-fold) compared to the controls (Fig. 6b). In line with the previous findings in ciPTEC, treatment of *CTNS^Patient^* tubuloids with the safe concentration of both cysteamine and bicalutamide (Supplementary fig. S5b-d) resulted in a more potent reduction in cystine levels (~2-fold) than with cysteamine treatment alone (Fig. 6c, d). Furthermore, we performed targeted metabolomic profiling (Supplementary file 2) and identified that bicalutamide (but not cysteamine), either alone or in combination with cysteamine, reduced the αKG levels in *CTNS^Patient-2^* tubuloids (Fig. 6e).

**Fig. 6.**
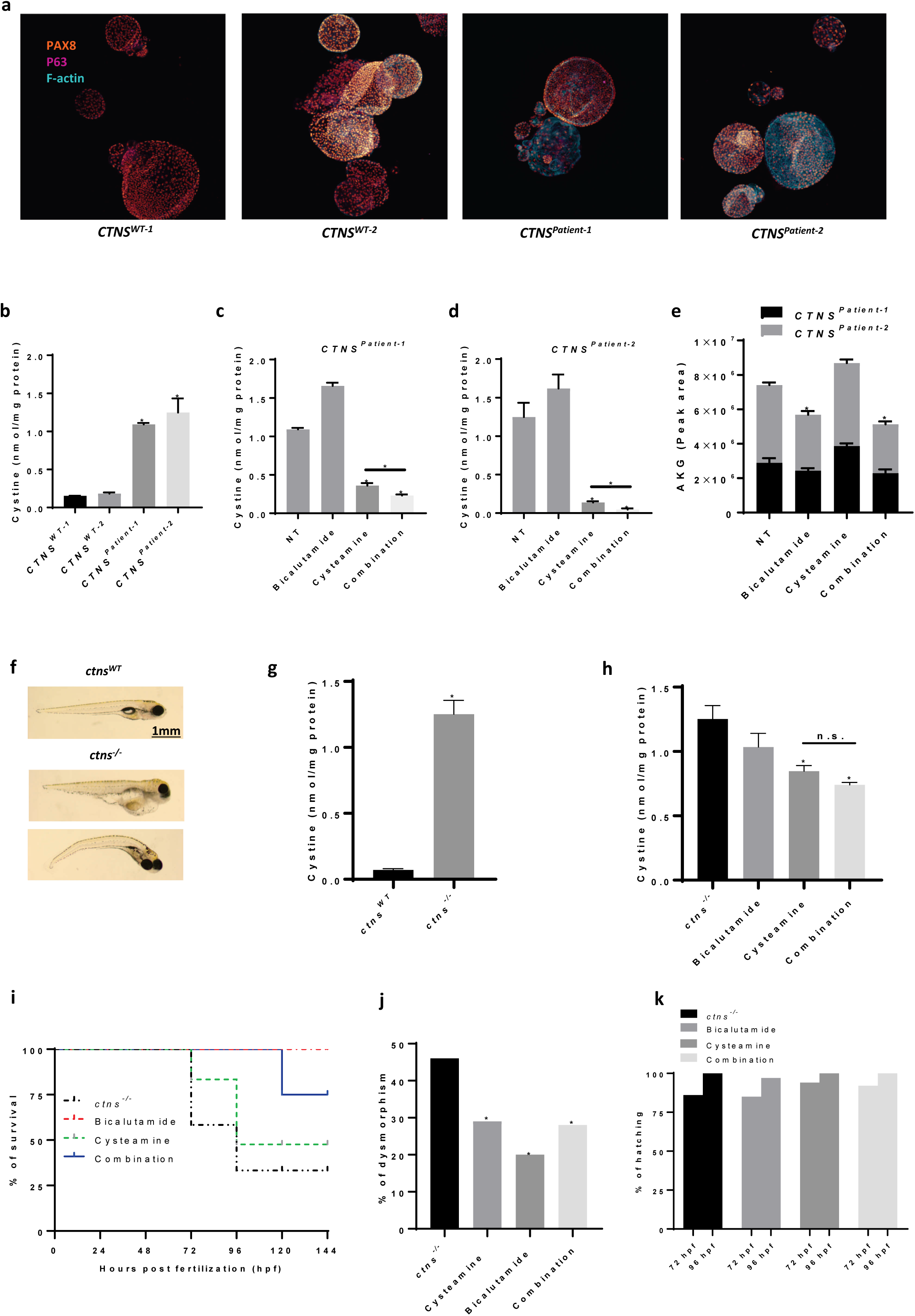
Cysteamine-bicalutamide combination treatment is effective in patient-derived cystinotic tubuloids and cystinotic zebrafish. **a**, Immunohistochemistry of patient-derived cystinotic tubuloids (*CNTS^Patient-1^* and *CTNS^Patient-2^*) and healthy kidney tissue-derived control tubuloids (*CNTS^WT-1^* and *CTNS^WT-2^*) for PAX8, p63 and F-actin. **b**, Quantification of cystine levels (nmol/mg protein) by HPLC-MS/MS in control and cystinotic tubuloids. **c**, **d**, Quantification of cystine levels (nmol/mg protein) by HPLC-MS/MS in two different patient-derived cystinotic tubuloids in the absence of the drug (NT) or upon treatment with cysteamine (100 μM), bicalutamide (35 μM) or cysteamine (100 μM)-bicalutamide (35 μM) combination treatment. **e**, αKG levels measured in patient-derived cystinotic tubuloids (*CNTS^Patient-1^* and *CTNS^Patient-2^*) in the absence of the drug (NT) or upon treatment with cysteamine (100 μM), bicalutamide (35 μM) or cysteamine (100 μM)-bicalutamide (35 μM) combination treatment using metabolomics. **f**, Representative images of control and cystinotic zebrafish. **g**, Quantification of cystine levels (nmol/mg protein) by HPLC-MS/MS in control and cystinotic zebrafish. **h**, Quantification of cystine levels (nmol/mg protein) by HPLC-MS/MS in cystinotic zebrafish after treatment with cysteamine (1000 µM), bicalutamide (10 µM), and a combination of cysteamine and bicalutamide (1000 µM and 10 µM, respectively). **i**, Survival rates in *ctns^−/−^* zebrafish upon treatment with cysteamine (1000 µM), bicalutamide (10 µM), and a combination of cysteamine and bicalutamide (1000 µM and 10 µM, respectively). **j**, Deformity rates in *ctns^−/−^* zebrafish after treatment with cysteamine (1000 µM), bicalutamide (10 µM), and a combination of cysteamine and bicalutamide (1000 µM and 10 µM, respectively). **k**, Hatching rates in surviving *ctns^−/−^* zebrafish evaluated at 72 and 96 hrs post fertilization (hpf) with cysteamine (1000 µM), bicalutamide (10 µM), and a combination of cysteamine and bicalutamide (1000 µM and 10 µM, respectively). The total numbers of embryos evaluated for survival, hatching, deformity rates, and cystine levels were 40 embryos per group. Drugs were administered at 48 hpf in all experiments dissolved in the swimming water with the specified concentrations. Data are expressed as the mean ± SEM of three independent experiments. Statistical analysis was performed using one way analysis of variance (ANOVA) followed by Dunnett’s multiple comparisons test. Where appropriate unpaired two-tailed Student’s t-test were used. P-values < 0.05 were considered to be significant.

### Bicalutamide and cysteamine combination treatment efficiently lowers cystine levels and improves survival of cystinotic zebrafish

Finally, we assessed the combination therapy in cystinotic zebrafish, a well-characterized *in vivo* model of cystinosis ^29,30^. Cystinotic zebrafish displayed significantly increased levels of cystine, dysmorphic features, delayed hatching, and reduced survival compared to the controls (Fig. 6f-k, supplementary fig. S6a-c). As expected, cysteamine was able to reduce cystine levels (1.5-fold; Fig. 6h) and there seemed to be an additive effect of bicalutamide (1.8-fold reduction for the combination) though not statistically significant. Further, bicalutamide either alone or in combination with cysteamine significantly improved the survival and reduce the percentage of dysmorphism in the cystinotic zebrafish without having an effect on their hatching rate, when compared to cysteamine monotherapy (Fig. 6i-k). This indicates the combination treatment is beneficial in cystinotic zebrafish without inducing toxicity.

## Discussion

In this work, we aimed to identify new therapeutic targets by evaluating the persistent cellular link between cystinosin loss-of-function and proximal tubular dysfunction in nephropathic cystinosis. Using an omics-based strategy we identified αKG as a key metabolite linking autophagy disruption, increased oxidative stress, and proximal tubule dysfunction in cystinotic renal proximal tubule cells. These data also indicated that a bicalutamide-cysteamine combination treatment could provide a novel pharmacological approach for the phenotypical correction of renal cystinosis.

Although several cellular defects have been associated with cystinosis, the exact mechanism linking cystinosin loss, lysosomal defects and epithelial dysfunction remains elusive. Defective cystinosin alters lysosomal autophagy dynamics ^31,32^, impairs activity of mTORC1 and increases the numbers of autophagosomes in *CTNS^−/−^* cells. Although our data suggest an abnormal induction of autophagy in cystinotic cells, the accumulation of the autophagy substrate SQSTM1 and the decreased lysosomal cargo processing, confirms the lack of autophagy completion in *CTNS^−/−^* cells, in agreement with the recent studies postulating an impairment of autophagy flux in many LSDs, including cystinosis ^33,34^. Evidence further supporting the defective degradative capacity of lysosomes in cystinotic proximal tubule cells include decreased lysosomal catalytic-proteins expression namely, lysosomal acid lipase, lysosomal acid phosphatase, lysosomal acid hydrolase, and cysteine proteases, cathepsin. Delivery of the newly synthesized lysosomal enzymes from Golgi to the lysosome relies mostly on the mannose 6-phosphate receptors ^35^, which was downregulated in *CTNS^−/−^* cells. We hypothesize that *CTNS* loss affects lysosomal function by hampering the delivery of newly synthesized lysosomal enzymes from Golgi to lysosomes and/or by inhibiting the maturation of these enzymes within (endo) lysosomal compartments. The latter hypothesis was confirmed by Festa *et al*. where they have shown the impairment of lysosome proteolysis (both *in vitro* and *in vivo*) is a result of defective cathepsin activation ^13^.

The question remains how cystinosin deficiency leads to autophagy dysfunction and epithelial impairment. By combining dual-omics and pharmacological approaches, here we show that there is a multifaceted impact of cystinosin loss-of-function in proximal tubule cells. Lysosomal accumulation of cystine leads to abnormal induction of autophagy providing more cargo to be digested in the lysosomes. This might lead to increased fusion of autophagosomes with cystine-loaded lysosomes, rendering them to be more susceptible to membrane destabilization and dysfunctional, ROS-producing mitochondria. Abnormal induction of autophagy and increased oxidative stress in parallel, results in rewiring of amino acid metabolism and fueling multiple metabolic pathways in *CTNS* deficient renal proximal tubule cells ^36,37^. This is in agreement with a recent study providing evidence that addition of amino acids, particularly cysteine, suppresses mTORC1 signaling and maintains autophagy in Drosophila larvae ^38^. The pathway involves the catabolism of cysteine to acetyl-CoA and subsequent fueling the TCA cycle intermediates, namely αKG. This confirms our finding where accumulation of αKG, a downstream metabolite of cysteine catabolism, was found to induce autophagy in cystinotic proximal tubule cells.

Another pathway that explains the over-production of αKG is the oxidative stress status in cystinosis. The link between cystinosin loss-of-function and increased oxidative stress have been long-established ^10,11,22,23^. αKG is a well-known antioxidant and its level can be increased by cells in response to oxidative stress contributing to the prevention and/or treatment of several disorders induced by oxidative stress ^19–21^. This is done through downregulation of the mitochondrial enzyme AKGDH, reported to be severely diminished in human pathologies where oxidative stress is thought to play a vital role ^21^. αKG together with cysteine and glycine, a is involved in GSH synthesis ^39,40^. GSH is one of the major cellular antioxidants, protecting cells against ROS. In cystinosis, however, due to dysfunctional cystinosin, both cystine and cysteine are trapped inside lysosomes and are less available for GSH synthesis, resulting in accumulation of αKG in cystinosis. The unbalanced production of αKG hence, abnormally induces autophagy and drives cystinotic cells into apoptosis.

Up till now, contradictory effects of αKG on autophagy have been reported. Several studies demonstrate that addition of the αKG to mammalian cells or its administration into mice effectively inhibits autophagy ^41–44^. In sharp contrast, Chin *et al*. have shown that αKG inhibits mTORC1 and the mitochondrial ATP synthase and therefore induces autophagy both in mammalian cells and in *Caenorhabditis elegans* ^26,45^. In agreement with the work of Chin *et al*., we showed that addition of the αKG precursor, DMKG, to cystinotic proximal tubule cells induces autophagy, demonstrated by increased LC3-II/LC3-I ratio.

Defective receptor mediated endocytosis, as a manifestation of renal Fanconi syndrome, is another hallmark of cystinosis ^32,46^. It is suggested that the cystinosin deficiency affects the expression of multiligand surface receptors, resulting in a delayed endocytotic cargo processing and reduced protein reabsorption in cystinotic patient-derived proximal tubule cells ^32^. CRISPR-mediated *CTNS* knockout proximal tubule cells, similar to cystinotic patient-derived cells, showed a delayed lysosomal cargo degradation but maintained their endocytosis function, postulating that the proximal tubule reabsorption machinery may not be primarily regulated by cystinosin and/or it could be a secondary consequence of cystinosis. The patient cells that have been most extensively studied in literature and have been used as a reference for this study (ciPTEC *CTNS^Patient^*) contain a 57kb genomic deletion of *CTNS*, sedoheptulose kinase (*SHPK*, also known as *CARKL*) and part of the potential vanilloid 1 (*TRPV1*) gene ^46,47^. *CARKL* and *TRPV1* genes loss in addition to *CTNS* may be linked to the reduced protein uptake seen in these cells. Moreover, culture adaptations and (epi)genetic variations between cell lines and donors can account for these variations.

To extend our knowledge on the beneficial effect of the combination treatment and bring the therapy one step closer to clinical application, we evaluated the safety and efficacy in patient-derived tubuloids, an advanced *in vitro* model, and *in vivo* in cystinotic zebrafish. Tubuloids offer several advantages over conventional mammalian cell lines. (1) They are human-derived three dimensional (3D) renal tubular epithelial cultures, offering physiological heterogeneity and recapitulating the *in vivo* situation. (2) Tubuloids are grown in 3D, allowing neighboring cells to interact in a more physiological way than in conventional 2D culture models. (3) Tubuloids are not genetically modified or reprogrammed and accurately mimic patients genotype and phenotype, allowing to develop a personalized medicine ^28^. The *ctns^−/−^* zebrafish, on the other hand, is a robust and versatile model of cystinosis with early phenotypic characteristics of the disease that can be used for the *in vivo* screening of novel therapeutic agents ^29,30^. The fact that the improved cystine lowering-efficacy of the combination treatment could be reproduced in tubuloids from two different patients and in cystinotic zebrafish, underlines the robustness of these findings and increases the likelihood that this treatment can be successfully extrapolated to cystinosis patients in general.

Autophagy blockade is one of the key features in LSDs, including cystinosis ^8,33^. Recent studies showing that overexpression of TFEB activity stimulates lysosomal excretion and rescues the delayed endocytic cargo processing, allowing the relief and/or treatment of the symptoms of many LSDs ^34^. This in line with the study of Rega *et al*. where stimulation of endogenous TFEB activity by genistein was shown to lower cystine levels and rescue the delayed endocytic cargo processing in cystinotic proximal tubule cells ^16^. Furthermore, inhibition of mTOR signaling by everolimus was shown to activate autophagy, rescue the number of large lysosomes, and in combination with cysteamine, reverse the cystine/cysteine loading defect in patient-specific and CRISPR-edited cystinotic induced pluripotent stem cells and kidney organoids ^48^.

Bicalutamide, a non-steroidal anti-androgenic agent, is one of the most widely prescribed drugs for treating prostate cancer ^49^. Recently, it has been patented for the treatment of several LSDs, promoting mTOR-associated autophagy and cellular exocytosis ^27^. Bicalutamide was able to increase endogenous TFEB activity, restore endocytic cargo processing, and in combination with cysteamine, efficiently normalize lysosomal cystine/cysteine levels in both patient-derived and CRISPR-edited cystinotic proximal tubule cells. Furthermore, cysteamine-bicalutamide combination treatment reversed the proteome and metabolic phenotype, especially αKG (through an upregulation of AKGDH), and resolved the αKG-mediated downstream effects in cystinotic proximal tubular cells. Although cysteamine alone could rescue cystine/cysteine loading defect and improve GSH levels, the main proposed mechanisms mediating αKG accumulation, it could not revert αKG levels in cystinotic cells. This phenomenon can be explained by the fact that cysteamine cannot correct the lysosomal defect in cystinosis.

Recently, it has been shown that cystinosin loss-of-function compromises autophagy-mediated clearance of dysfunctional mitochondria, which are the potent source of ROS-producing compounds ^13^. In this context, the αKG lowering effect of bicalutamide can be explained by its autophagy-activating properties, where it stimulates cellular exocytosis and restores endocytic cargo processing, leading to a decreased autophagy-mediated ROS production in cystinosis. Moreover, bicalutamide combined with cysteamine can efficiently replete the cytoplasmic pool of cystine/cysteine from the lysosomal compartment, making them more available for αKG to produce GSH, further confirming the importance of the combination therapy in cystinosis.

It is worth mentioning that, the use of any androgen deprivation therapy are generally associated with the risk of acute kidney injury (AKI) ^50,51^. Up to 36% of patients who have taken bicalutamide for 1 to 6 months may experience AKI ^52^. However, this association is mainly driven by a combination of gonadotropin- or luteinizing-releasing hormone agonists with bicalutamide ^50^. Bicalutamide-cysteamine combination treatment at a concentration to have biological activity was not only found to be safe in our models, but also improved the survival and reduce the percentage of dysmorphism in the cystinotic zebrafish, confirming the safety profile of bicalutamide both in *in vitro* and *in vivo*. Anti-androgenic therapy is also known to reduce testosterone levels ^50,53^, complicating the beneficial effect of bicalutamide in cystinotic male patients with delayed maturation ^54–56^. Bicalutamide is a racemate and its anti-androgenic activity resides solely in the (R)-enantiomer ^57,58^, while the (S)-enantiomer with little, if any, anti-androgenic activity mediates autophagy effects ^27^. Therefore, it would be of great scientific benefit to further investigate the effect of the (S)-bicalutamide and/or develop any structural analogues to treat cystinosis.

Taken together, we identified a new therapeutic target by evaluating the persistent cellular abnormalities using an omics-based strategy. We identified αKG as a key metabolite linking cystinosin loss, lysosomal autophagy defect, and proximal tubular impairment in cystinosis. Bicalutamide, but not cysteamine, was able to normalize αKG levels and resolve αKG-mediated downstream effects in cystinotic cells. Bicalutamide in combination with cysteamine demonstrated an additive effect and resulted in a more potent reduction in cystine levels than with cysteamine treatment both in *in vitro* and *in vivo*, suggesting this combination therapy may hold great potential to treat patients with cystinosis.

## Methods

### Reagent and antibodies

All chemicals and reagents were obtained from Sigma-Aldrich (Zwijndrecht, The Netherlands) unless specified otherwise. Primary antibodies used were mouse anti-LAMP1 (Santa Cruz Biotechnology #sc-18821, dilution 1:200), rabbit anti-mTOR (Cell Signaling Technology #2983, dilution 1:400), rabbit anti-LC3 (Novus Biologicals #NB600-1384SS, dilution 1:1000), mouse anti-SQSMT1 (p62) (BD Biosciences #610832, dilution 1:1000), rabbit anti-β-actin (13E5) (Cell signaling technology #4970, dilution 1:4000), Rabbit anti-PAX8 (Proteintech, dilution 1:200), mouse anti-p63 (Abcam, dilution 1:200), and Phalloidin-AF488 (Life Technology, dilution 1:100). Polyclonal goat anti-rabbit (#P0448, dilution 1:5000) and polyclonal goat anti-mouse (#P0447, dilution 1:5000) secondary antibodies were obtained from Dako products (CA, USA). Alexa-488 goat anti-mouse (dilution 1:500), Alexa-647 goat anti-rabbit (dilution 1:200), donkey anti-rabbit-AF647 (dilution 1:300), and donkey anti-mouse-AF568 (dilution 1:200) secondary antibodies were from Life Technologies Europe BV (The Netherlands).

### Generation of *CTNS^−/−^* isogenic cell line of ciPTEC using CRISPR

Guide RNAs (gRNAs) targeting exon 4 of the *CTNS* gene were designed using the online gRNA designing tool available at chopchop.cbu.uib.no. In order to maximize specificity, guide sequences with high scores for on-target efficiency and no predicted off-targets having at least 3 base pair mismatches in the genome were selected. Optimal gRNA (5’-GTCGTAAAGCTGGAGAACGG-3’) was cloned into the pSPCas9(BB)-2A-GFP plasmid (Addgene #48138) as described previously by Ran et al. ^59^, and introduced into healthy ciPTEC using PolyPlus JetPrime. 72 hrs post-transfection, GFP-positive singlet cells were sorted using FACS Aria-II flow cytometer and expanded in 96-wells plate. The gRNA cut site was amplified with PCR using the primers flanking the cut region (F.CTNS_ex4 5’-GGCCTGTTTTCCTCCATCTCTG-3’; R.CTNS_ex4 5’-AAGTGCCAACCAGCAGCTC-3’). knockouts were confirmed by Sanger sequencing and subsequently by intracellular cystine accumulation.

### CiPTEC culture

The ciPTEC were cultured as described previously by Wilmer et al. 2010 ^60^. The culture medium was Dulbecco’s modified Eagle medium DMEM/F-12 (GIBCO) supplemented with fetal calf serum 10% (v/v), insulin 5 µg/ml, transferrin 5 µg/ml, selenium 5 µg/ml, hydrocortisone 35 ng/ml, epidermal growth factor 10 ng/ml and tri-iodothyronine 40 pg/ml. In short, cells were seeded at a density of 55,000 cells/cm^2^ and grown at 33°C for 24 hrs to enable them to proliferate and subsequently cultured at 37°C for 7 days to mature into fully differentiated epithelial cells. As a reference we also included a non-isogenic patient derived *CTNS^−/−^* ciPTEC line, which will be referred to as *CTNS^Patient^*.

### Tubuloid culture

Tubuloids were established from the urine of two pediatric cystinosis patients and healthy kidney tissue from two donors as previously described ^28^. Urine was collected, cooled to 4°C, rinsed with phosphate buffered saline (PBS) and with advanced Dulbecco’s modified Eagle’s medium (ADMEM)-F12 (Gibco) supplemented with 1% HEPES, 1% glutamax, 1% penicillin/streptomycin, 0,1 mg/mL primocin (Invivogen) and 10 μM Y-27632 (Abmole). The pellets were resuspended in basement membrane extract (BME, SanBio/Trevigen) and plated. After BME droplets had solidified, ADMEM-F12 supplemented with 1% HEPES, 1% glutamax, 1% penicillin/streptomycin, 0,1 mg/mL primocin, 1,6% B27 (Gibco), 1% Rpsondin-3 conditioned medium (U-Protein Express), 50 ng/mL EGF (Peprotech), 100 ng/mL FGF10 (Peprotech), 1 mM N-acetylcysteine, 5 μM A83-01 (Tocris Bioscience) and 10 μM Y-27632 was added as “expansion medium”. In order to establish tubuloids from kidney tissue, the tissue was digested with 1 mg/mL collagenase for 45 minutes. Tissue fragments were resuspended in BME and plated. After BME droplets had solidified, expansion medium was added. For specific experiments, tubuloids were differentiated using ADMEM-F12 with 1% HEPES, 1% glutamax and 1% penicillin/streptomycin (differentiation medium).

### Cell treatment

Standard starvation medium was Hank’s balanced salt solution (HBSS; GIBCO). When indicated, the cell-permeable form of α-ketoglutarate, dimethyl α-ketoglutarate (DMKG) was added to a final concentration of 2 mM for 4 or 24 hrs. Cysteamine (100 μM), bicalutamide (35 μM), or the combination of cysteamine and bicalutamide (100 μM and 35 μM, respectively) were used as a treatment for 96 hrs (5 days).

### Zebrafish maintenance and breeding

The animal care and experimental procedures were carried out in accordance with the ethical committee guidelines for laboratory animal experimentation at KU Leuven. Zebrafish (Danio rerio) were AB strain wild-type and *ctns^−/−^* mutant. Cystinotic and wild-type larvae were raised at 28.5 °C in egg water (Instant Ocean Sea Salts, 60μg/ml). At 48 hrs post fertilization, larvae were treated with cysteamine 1000 µM, bicalutamide 10 µM, or their combination. Drugs were administered at 48 hrs post fertilization in all experiments dissolved in the swimming water. The medium was refreshed every day and dead embryos were sorted out.

### Intracellular cystine quantification by HPLC-MS/MS

Cystine levels were quantified using HPLC-MS/MS; a rapid and sensitive assay that has been developed and validated in house ^61^. In brief, ciPTEC, tubuloids, and zebrafish larvae pellets were suspended in N-Ethylmaleimide (NEM) solution containing 5 mM NEM in 0.1 mM sodium phosphate buffer (pH 7.4). The cell suspension were then precipitated and protein was extracted with sulfosalicylic acid 15% (w/v) and centrifuged at 20,000 g for 10 min at 4°C. Protein concentration was determined by the method of the Pierce™ BCA protein assay kit according to the manufacturer’s protocol (Thermo Fischer, The Netherlands), and the cystine concentration was measured using HPLC-MS/MS. Data are expressed as the cystine values (nmol) corrected for total protein content (mg).

### Quantitative real-time PCR

The mRNAs were extracted from cells using the Qiagen RNeasy mini kit according to the manufacturer’s instructions. Total mRNA (600 ng) was reverse transcribed using iScript Reverse Transcriptase Supermix (Bio-Rad). Quantitative real-time PCR was performed using iQ Universal SYBR Green Supermix (Bio-Rad) with the specific sense and anti-sense primers for *TFEB* (forward: 5’-GCAGTCCTACCTGGAGAATC-3’; reverse: 5’-GTGGGCAGCAAACTTGTTCC-3’). The ribosomal protein S13 (*RPS-13*) (forward: 5’-GCTCTCCTTTCGTTGCCTGA-3’; reverse: 5’-ACTTCAACCAAGTGGGGACG-3’) was used as the reference gene for normalization and relative expression level were calculated as fold change using the *2^−ΔΔ^Ct* method.

Tubuloids were grown in expansion medium for a few days and then differentiated for 7 days. Tubuloids were lysed and RNA was isolated using the RNEasy Mini Kit (Qiagen) according to the manufacturer’s protocol. Quantitative real-time qPCR was performed using the iQ SYBR Green Supermix (Bio-Rad). A 384-wells plate was used with a reaction volume of 12,5 μL and duplicates for each reaction. For the read-out, the CFX384 Touch Real-Time PCR Detection System (Bio-Rad) was used. Expression levels of the following genes were measured: *RPS-13* (housekeeping gene), *ANPEP*, *ABCC3*, *HNF1A*, *HNF4A*, *SLC12A1*, *SLC12A3*, *CALB1*, *AQP2* and *AQP3*. Primer sequences are provided in supplementary Table. 1. Expression was normalized to the expression of *RPS-13* within the same sample (ΔCt).

### Immunofluorescence and confocal microscopy

To investigate mTORC1/LAMP1 co-localization, control and cystinotic ciPTEC lines were cultured on coverslips with the respective treatments. Thereafter, cells were fixed with 4% paraformaldehyde in PBS for 10 min, permeabilized with 0.1% Triton-X solution for 10 min, and blocked with 1% bovine serum albumin (BSA) diluted in PBS for 30 min. Subsequently, cells were stained with the primary antibodies diluted in blocking buffer overnight at 4°C. After 3 washes with PBS, the cells were incubated for 2 hrs at room temperature with the corresponding secondary antibodies. Nuclei were stained with Hoechst 33342 (1 µM) and cells were imaged with a Deltavision confocal microscope (Cell Microscopy Core, Department of Cell Biology, University Medical Centre Utrecht).

To assess TFEB intracellular distribution, cells were seeded in special optic 96-wells plate until to reach 50% confluence. Cells were then transfected with the TFEB-GFP plasmid (a kind gift from Dr. Annelies Michiels (Viral Vector Core, Leuven, Belgium)) using PolyPlus JetPrime reagent according to the manufacturer’s instructions. After 48 hrs from transient transfection, cells were stained with Hoechst 33342 (1 µM) for 10 min and imaged using a Cell Voyager 7000 (CV7000) confocal microscope (Yokogawa Electric corporation, Tokyo, Japan). TFEB nuclear translocation data are expressed as number of cells with nucleus-TFEB positive over the total number of TFEB-transfected cells.

For tubuloid experiments, tubuloids were differentiated for 7 days. Tubuloids were fixed for 45 min with 4% formaldehyde in PBS and then permeabilized and blocked with 0.5% Triton-X100 plus 0,5% BSA in PBS for 30 min. Subsequently, tubuloids were stained with the primary antibodies diluted in 0.5% BSA plus 0.1% Tween20 in PBS overnight at 4°C. Tubuloids were rinsed twice with 0.5% BSA plus 0.1% Tween20 in PBS and then stained for 2 hrs at room temperature with the corresponding secondary antibodies and DAPI (1:1000) in 0.5% BSA plus 0.1% Tween20 in PBS. Tubuloids were washed twice with 0.5% BSA + 0.1% Tween20 in PBS, mounted and imaged using a Leica SP8 confocal microscope.

### Immunoblots

The ciPTEC were seeded in 6-well plates in triplicate with the respective treatments. Subsequently, cells were washed twice with PBS and lysed with RIPA buffer containing protease inhibitor cocktail (Roche). Protein quantification was performed using the method of the Pierce™ BCA protein assay kit according to the manufacturer’s protocol. After the electrophoresis, the proteins were transferred to a nitrocellulose membrane (midi kit, Bio-Rad) with Trans-Blot Turbo Transfer System (Bio-Rad). Finally, membranes were imaged using the ChemiDoc™ XRS+ (Bio-Rad) and analyzed using image J software.

### Endocytosis assay

The endocytic uptake was monitored in ciPTEC following incubation for 1.5 hrs at 37°C with 50 μg/ml of either BSA-AlexaFluor-647 (A34785, Thermo Fisher Scientific) or DQ Red BSA (D12051, Invitrogen). The cells were then fixed and stained with Hoechst 33342 (1 µM) for 10 min and imaged using a CV7000 confocal microscope (Yokogawa Electric corporation, Tokyo, Japan). Data were quantified with Columbus™ Image Data Storage and analysis software (PerkinElmer, Groningen, The Netherlands). Data are expressed as the number of BSA/DQ-BSA spots per cell.

### UHPLC/MS system and analysis of metabolomics data

#### Sample preparation

CiPTEC cells were washed with ice cold PBS and metabolites were extracted in 1 mL lysis buffer containing methanol/acetonitrile/dH_2_O (2:2:1). Samples were centrifuged at 16.000g for 15 minutes at 4 °C and supernatants were collected for LC-MS analysis.

#### Mass spectrometry and data analysis

LC-MS analysis was performed on an Exactive mass spectrometer (Thermo Scientific) coupled to a Dionex Ultimate 3000 autosampler and pump (Thermo Scientific). The MS operated in polarity-switching mode with spray voltages of 4.5 kV and -3.5 kV. Metabolites were separated using a Sequant ZIC-pHILIC column (2.1 x 150 mm, 5 μm, guard column 2.1 x 20 mm, 5 μm; Merck) with elution buffers acetonitrile (A) and eluent B (20 mM (NH4)2CO3, 0.1% NH4OH in ULC/MS grade water (Biosolve)). Gradient ran from 20% eluent B to 60% eluent B in 20 minutes, followed by a wash step at 80% and equilibration at 20%, with a flow rate of 150 μl/min. Analysis was performed using LCquan software (Thermo Scientific). Metabolites were identified and quantified on the basis of exact mass within 5 ppm and further validated by concordance with retention times of standards. Peak intensities were normalized based on total peak intensities and data were analyzed using MetaboAnalyst ^62^.

### UHPLC/MS-MS system and analysis of proteomics data

#### Sample preparation

CiPTEC pellets were lysed in boiling guanidinium lysis buffer containing 6 M guanidinium HCl (GuHCl), 5 mM tris(2-carboxyethyl)phosphine (TCEP), 10 mM chloroacetamide, 100 mM Tris-HCl pH 8.5, supplemented with protease inhibitor (cOmplete mini EDTA-free, Roche). Pellets were boiled for 10 min at 99 °C, sonicated for 12 rounds of 5 seconds (Bioruptor Plus, Diagenode), and spun down at 20,000 x g for 15 min. Protein concentration was determined using Pierce™ BCA protein assay kit. Equal amounts of protein per condition were digested with Lys-C (1:100, Wako) for 4 hrs at 37 °C, diluted to a final concentration of 2 M GuHCl, followed by trypsin digestion (1:100, Sigma Aldrich) overnight at 37 °C. Tryptic peptides were acidified to a final concentration of 1% formic acid (FA) (Merck), cleaned up using SepPak cartridges (Waters), and dried *in vacuo*.

#### Mass spectrometry and data analysis

Peptide samples were analyzed with an UHPLC 1290 system (Agilent technologies) coupled to an Orbitrap Q Exactive HF X mass spectrometer (Thermo Scientific). Peptides were trapped (Dr Maisch Reprosil C18, 3 μm, 2 cm x 100 μm) and then separated on an analytical column (Agilent Poroshell EC-C18, 2.7 μm, 50 cm x 75 μm). Trapping was performed for 5 min in solvent A (0.1% FA) and eluted with following gradient: 4 - 8% solvent B (0.1% FA in 80% acetonitrile) in 4 min, 8 - 24% in 158 min, 24 - 35% in 35 min, 35 - 60% in 17 min, 60 - 100% in 4 min and finally 100 % for 1 min. Flow was passively split to 300 nl/min. The mass spectrometer was operated in data-dependent mode. At a resolution of 35.000 *m/z* at 400 *m/z*, MS full scan spectra were acquired from *m/z* 375–1600 after accumulation to a target value of 3e^6^. Up to 15 most intense precursor ions were selected for fragmentation. HCD fragmentation was performed at normalised collision energy of 27% after the accumulation to a target value of 1e^5^. MS/MS was acquired at a resolution of 30,000. Dynamic exclusion was enabled with an exclusion duration of 32s. RAW data files were processed with MaxQuant (v1.6.0.16 (Cox, Mann 2008)) and MS2 spectra were searched with the Andromeda search engine against the Swissprot protein database of Homo Sapiens (20,259 entries, downloaded 31/01/2018) spiked with common contaminants. Cysteine carbamidomethylation was set as a fixed modification and methionine oxidation and protein N-term acetylation were set as variable modifications. Trypsin was specified as enzyme and up to two miss cleavages were allowed. Filtering was done at 1% false discovery rate (FDR) at the protein and peptide level. Label-free quantification (LFQ) was performed, and “match between runs” was enabled. The data was further processed using Perseus 1.6.0.7 ^63^.

#### Alpha-ketoglutarate measurement by LC-MS/MS

Alpha-ketoglutarate (2-KG) (fluka 75893) calibration standards of respectively 0.8, 1.6, 4.0, 8.0, and 20 µM, and an internal standard solution of 2.2 µM 3,3,4,4-D4 2kg (kind gift Prof. Dr. H. Blom VU Amsterdam) were prepared in dH2O. 50 µl plasma sample or calibration standard together with 50 µl internal standard solution and 50 µl dH2O were pipetted onto a Microcon ultrafilter (30 kDa, Millipore) The resulting ultrafiltrate (15min 14000g 15°C) was acidified with 20 µl 4% formic acid (Merck) in dH2O. 3 µl of the acidified ultra-filtrate was injected onto a peek-lined InertSustain AQ-c18 (2.1*100mm dp 3µ) column using a I-Class Acquity (Waters) The column was run at 40°C in gradient mode using 0.5% acetic acid in H2O and Acetonitrile (initial conditions 100% 0.5% acetic acid in H2O at 250 µl/min). The column flow was directed to a Xevo TQSµ (Waters) fitted with an electrospray ionization probe operating in the negative mode at unit resolution. The capillary voltage was set at 0.6 kV. The temperature settings for the source and ionblock were respectively 550°C and 150°C. As a drying gas nitrogen was used at a flow rate of 800 L/h. The conegas flow was set at 50 L/h. The collision cell was operated with argon as the collision gas at a pressure of 0.35 Pa. An area response was generated by recording the MRM transitions of the neutral loss CO2 for both 2kg an d4-2kg. The d4-2kg was used to normalize the area response. Quantification was performed using a linear regression curve constructed from the normalized area response from the prepared calibration standards.

### Cell viability assays

To estimate the percentage of cell death, cells were seeded in a 96-well plate and after respective treatments, the percentage cell death was assessed using Presto Blue Cell Viability Reagent according to manufacturer’s instructions.

For tubuloid experiments, the drug safety screening was performed using a protocol derived from Driehuis et al ^64^. In short, BME was dissolved by incubation in 1 mg/mL dispase II for 30 min at 37 °C. Tubuloids were digested into tiny fragments using Accutase. These fragments were plated in BME, grown for 1 day in expansion medium and then switched to differentiation medium for 2 days. Next, BME was dissolved using dispase II as described above, after which tubuloids were sieved using a 70 μm cell strainer (BD Falcon) and counted. 500 tubuloids in 40 μL differentiation medium with 5% BME were plated in a 384-wells plate using the Multidrop dispenser (Thermofisher) and treated with increasing concentrations of bicalutamide that were always combined with 100 μM cysteamine. Staurosporin (10 μM) was used as positive control. DMSO volume was 1% in all conditions. Drugs were added using the D300E digital dispenser (Tecan). After 5 days, tubuloids were lysed with 40 μL Cell-Titer Glo (Promega). Luminescence was measured using the Spark® multimode microplate reader (Tecan) to determine ATP levels as an indicator of the amount of tubuloids that had survived the treatment. Tubuloid viability was normalized to the viability (ATP levels) upon treatment with cysteamine alone (=100%).

### Apoptosis assay

The ciPTEC were plated in a optic 96-well imaging plates and treated with DMKG in presence or absence of bicalutamide, cysteamine or their combination. Subsequently, the Cell Event Caspase- 3/7 Green Detection Reagent (8 µM) was added and incubated for 30 min before imaging. Caspase-active cells were identified as described in manufacturer’s instruction. Each well was imaged using the CV7000 confocal microscope and analyzed with Columbus™ Image Data Storage and analysis software (PerkinElmer, Groningen, The Netherlands).

### ROS detection assay

The ciPTEC were seeded in 96-well plates and ROS levels were assessed using general oxidative stress indicator (CM-H2DCFDA; Invitrogen) according to the manufacturer’s protocol. Briefly, the cells were treated with the CM-H2DCFDA reagent (10 μM) and incubated in the dark. Following 20 min incubation at 37°C, cells were rinsed once with HBSS and incubated with the different treatment conditions as previously stated. Fluorescence was measured using a fluorescent microplate reader (Fluoroskan Ascent, Thermo Fisher Scientific, Vantaa, Finland) at excitation wavelength of 492 nm and emission wavelength of 518 nm. Data are expressed as the fluorescence values normalized to the protein concentration.

### Statistical analysis

Statistical analysis was performed using GraphPad Prism 7.0 (GraphPad Software, Inc., USA). Data are presented as mean ± standard error of the mean (SEM) of minimally three independent experiments performed in triplicate. Significance was evaluated using One-way Analysis of Variance (ANOVA), or where appropriate unpaired two-tailed Student’s t-test was applied. P-values < 0.05 were considered to be significant.

### Data availability

The authors declare that all the data supporting the findings of this study are available within the article and its supplementary information files and from the corresponding author on reasonable request.

## Acknowledgements

This work was financially supported by a grant from the Dutch Kidney Foundation (grant nr.150KG19).

## Supplementary figures

**Supplementary figure S1.**
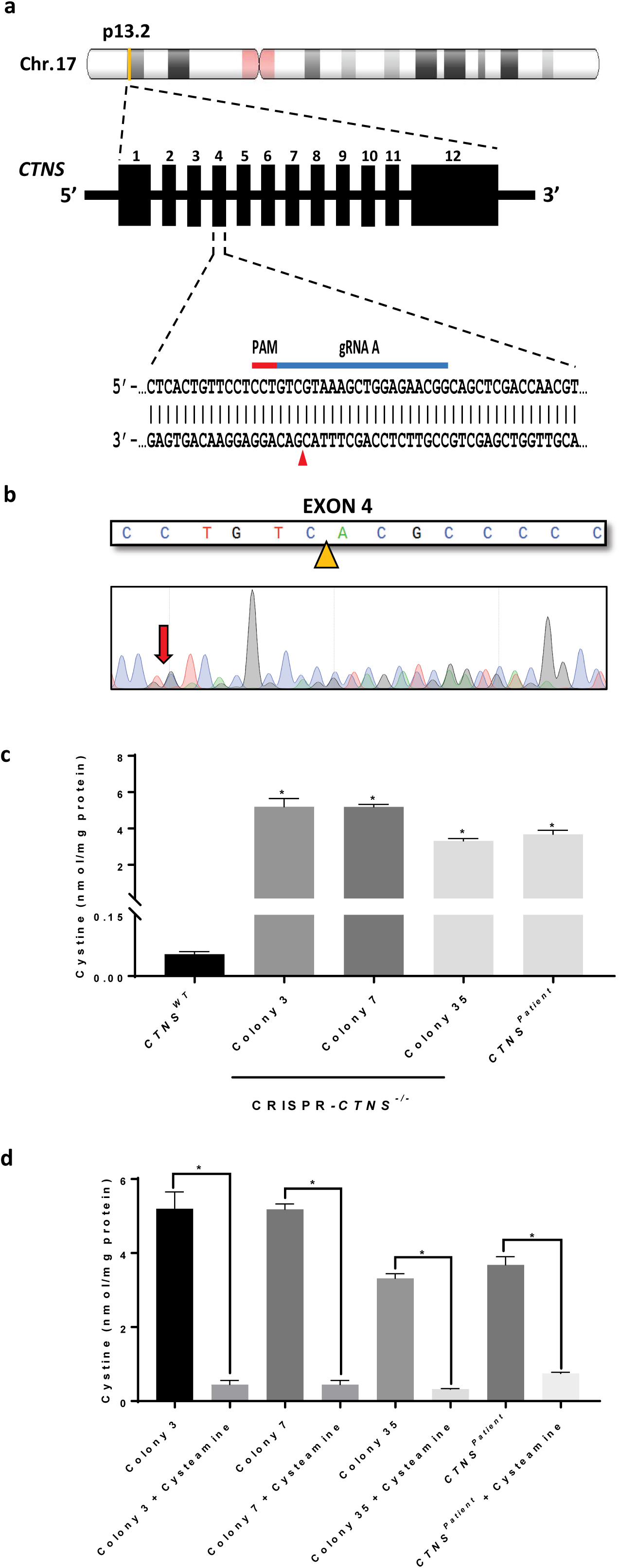
Generation of *CTNS^−/−^* isogenic cell line of ciPTEC using CRISPR. **a**, Schematic overview of the CRISPR-based strategy to knockout the *CTNS* gene in ciPTEC. **b**, Sanger sequencing chromatogram shows resulting sequence in CRISPR-generated cystinotic cells (*CTNS^−/−^*). **c,** Quantification of cystine levels (nmol/mg protein) by HPLC-MS/MS in in control (*CTNS^WT^*), CRISPR-generated cystinotic cells (*CTNS^−/−^*; line 3, 7, and 35), and patient-derived cystinotic cells (*CTNS^Patient^*). **d**, Quantification of cystine levels (nmol/mg protein) by HPLC-MS/MS in *CTNS^−/−^* lines (3, 7, and 35), and *CTNS^Patient^* cells upon treatment with cysteamine (100 µM). Data are expressed as the mean ± SEM of three independent experiments. Statistical analysis was performed using one way analysis of variance (ANOVA) followed by Dunnett’s multiple comparisons test. Where appropriate unpaired two-tailed Student’s t-test were used. P-values < 0.05 were considered to be significant.

**Supplementary figure S2.**
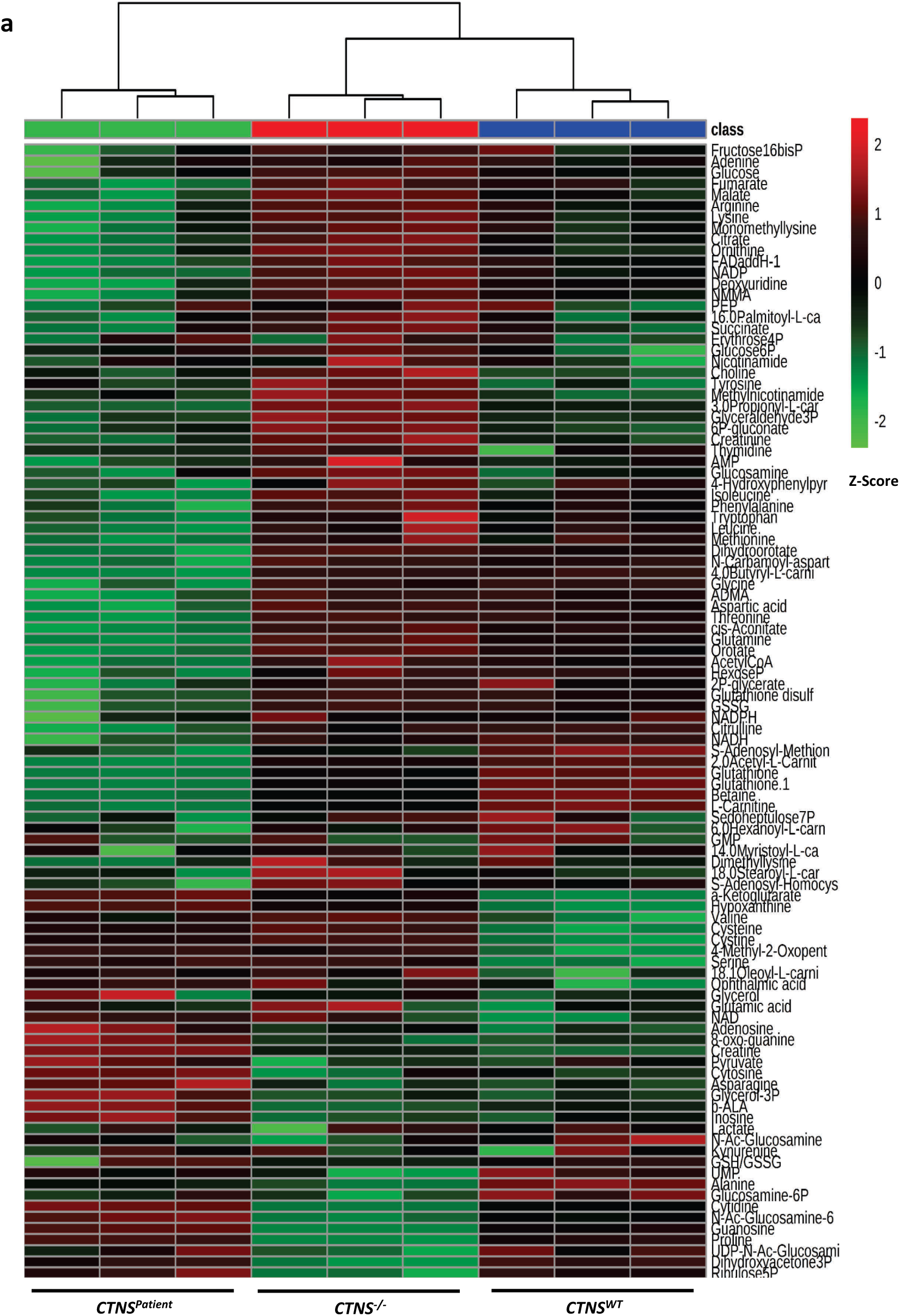
CRISPR-generated *CTNS^−/−^* cells are a good isogenic cystinosis model of healthy control cells. **a**, Heatmap analysis of metabolites distinctively expressed in control (*CTNS^WT^*), CRISPR-generated cystinotic cells (*CTNS^−/−^*), and patient-derived cystinotic cells (*CTNS^Patient^*). The row displays metabolite feature and the column represents the samples. The row Z-score of each feature is plotted in red-green color scale. Metabolites significantly decreased (P < 0.05) were displayed in green, while metabolites significantly increased (P < 0.05) were displayed in red.

**Supplementary figure S3.**
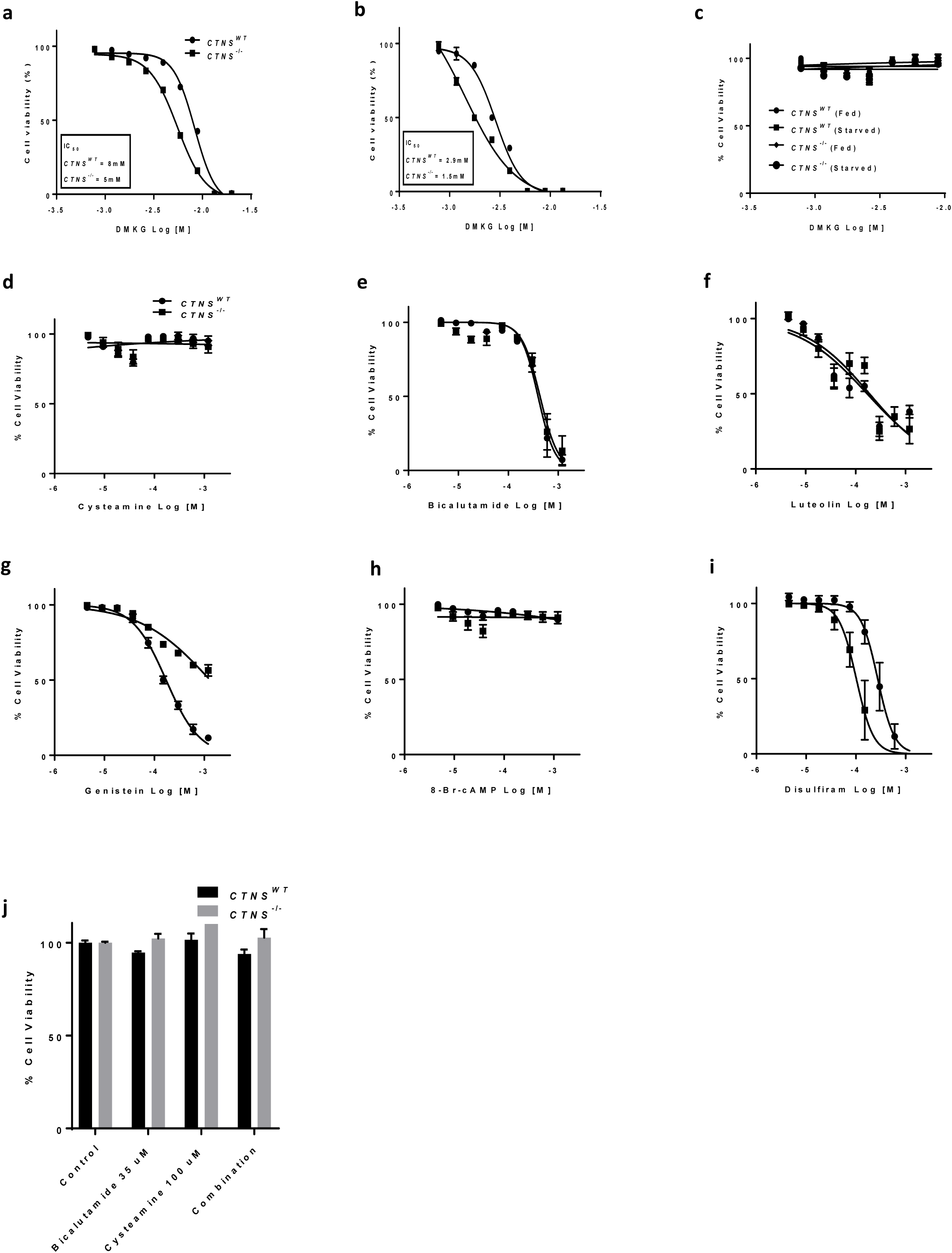
**a, b,** Cell viability curves of the increasing concentration of dimethyl α-ketoglutarate (DMKG) in control (*CTNS^WT^*) and CRISPR-generated cystinotic (*CTNS^−/−^*) cells after 24 hrs of incubation in fed and starved condition, respectively. **c**, Cell viability curves of DMKG in *CTNS^WT^* and *CTNS*^−/−^ cells after 4 hrs of incubation in fed and starved condition. **d**-**j**, Cell viability test in *CTNS^WT^*, *CTNS^−/−^* cells treated with the increasing concentrations of cysteamine, bicalutamide, luteolin, genistein, 8-bromo-cAMP, disulfiram, and a combination of cysteamine and bicalutamide (100 µM and 35 µM, respectively), respectively. Statistical analysis was performed using one way analysis of variance (ANOVA) followed by Dunnett’s multiple comparisons test. Where appropriate unpaired two-tailed Student’s t-test were used. P-values < 0.05 were considered to be significant.

**Supplementary figure S4.**
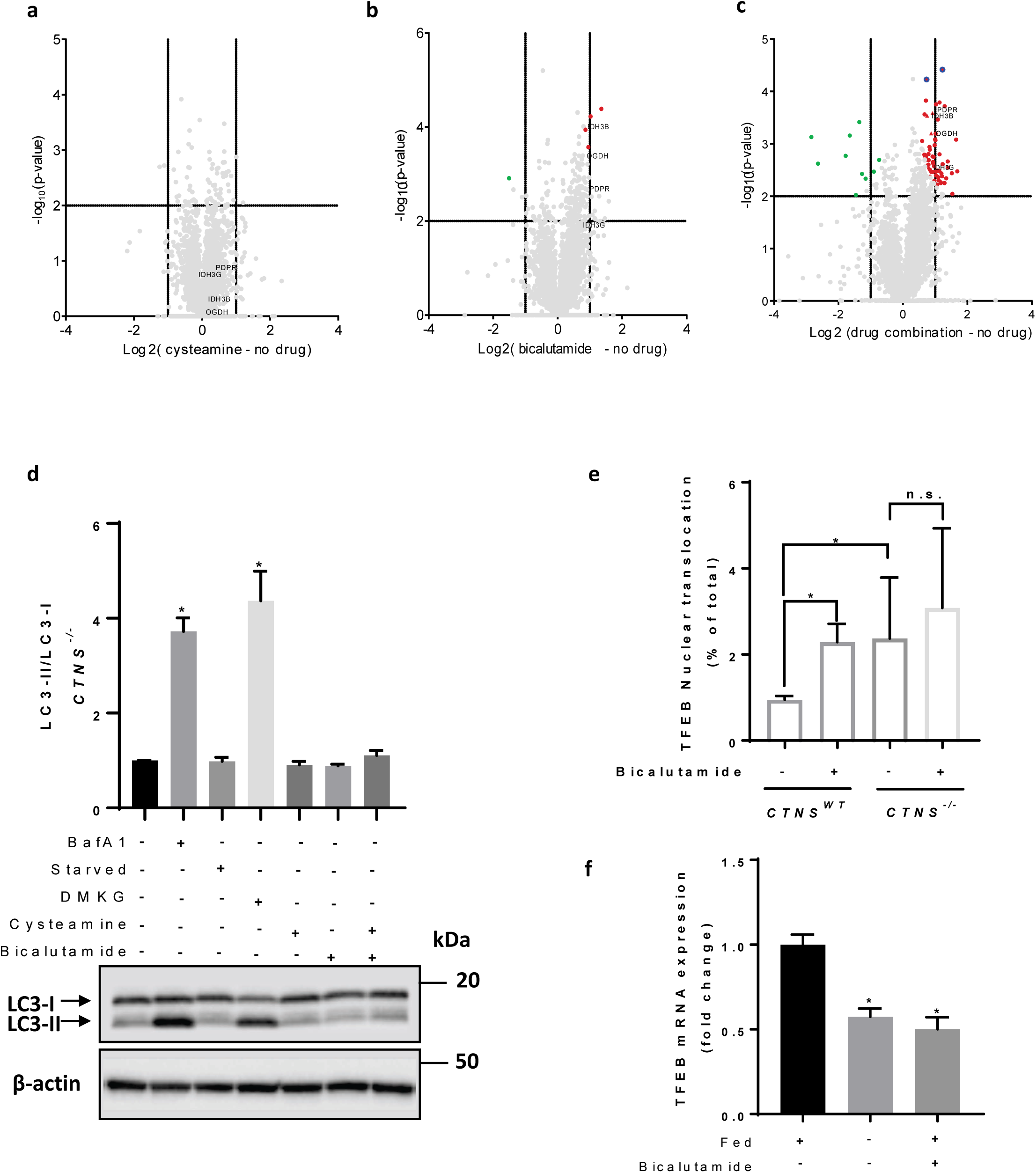
Cysteamine-bicalutamide combination treatment shows a synergic effect in treatment of cystinotic ciPTEC. **a**, **b**, **c**, Volcano plot illustrates significantly differentially abundant proteins. The -log_10_ (Benjamini–Hochberg corrected P-value) is plotted against the log_2_ (fold change: *CTNS^−/−^* no drug treatment/*CTNS^−/−^* cysteamine treatment), (fold change: *CTNS^−/−^* no drug treatment/*CTNS^−/−^* bicalutamide treatment), and (fold change: *CTNS^−/−^* no drug treatment/*CTNS^−/−^* combination treatment), respectively. The non-axial vertical lines denote ±1.5- fold change while the non-axial horizontal line denotes P = 0.05, which is our significance threshold (prior to logarithmic transformation). **d**, Western blotting and densitometric analyses for LC3-II/LC3-I ratio in CRISPR generated *CTNS^−/−^* cells treated with cysteamine (100 µM), bicalutamide (35 µM), and a combination of cysteamine and bicalutamide (100 µM and 35 µM, respectively). β-Actin was used as a loading control. **e**, Quantification of TFEB-GFP nuclear translocation in *CTNS^WT^*, and *CTNS^−/−^* cells upon treatment with bicalutamide (35 µM). **f**, *TFEB* mRNA expression of the *CTNS^WT^* cells upon starvation and treatment with bicalutamide (35 µM). Statistical analysis was performed using one way analysis of variance (ANOVA) followed by Dunnett’s multiple comparisons test. P-values < 0.05 were considered to be significant.

**Supplementary figure S5.**
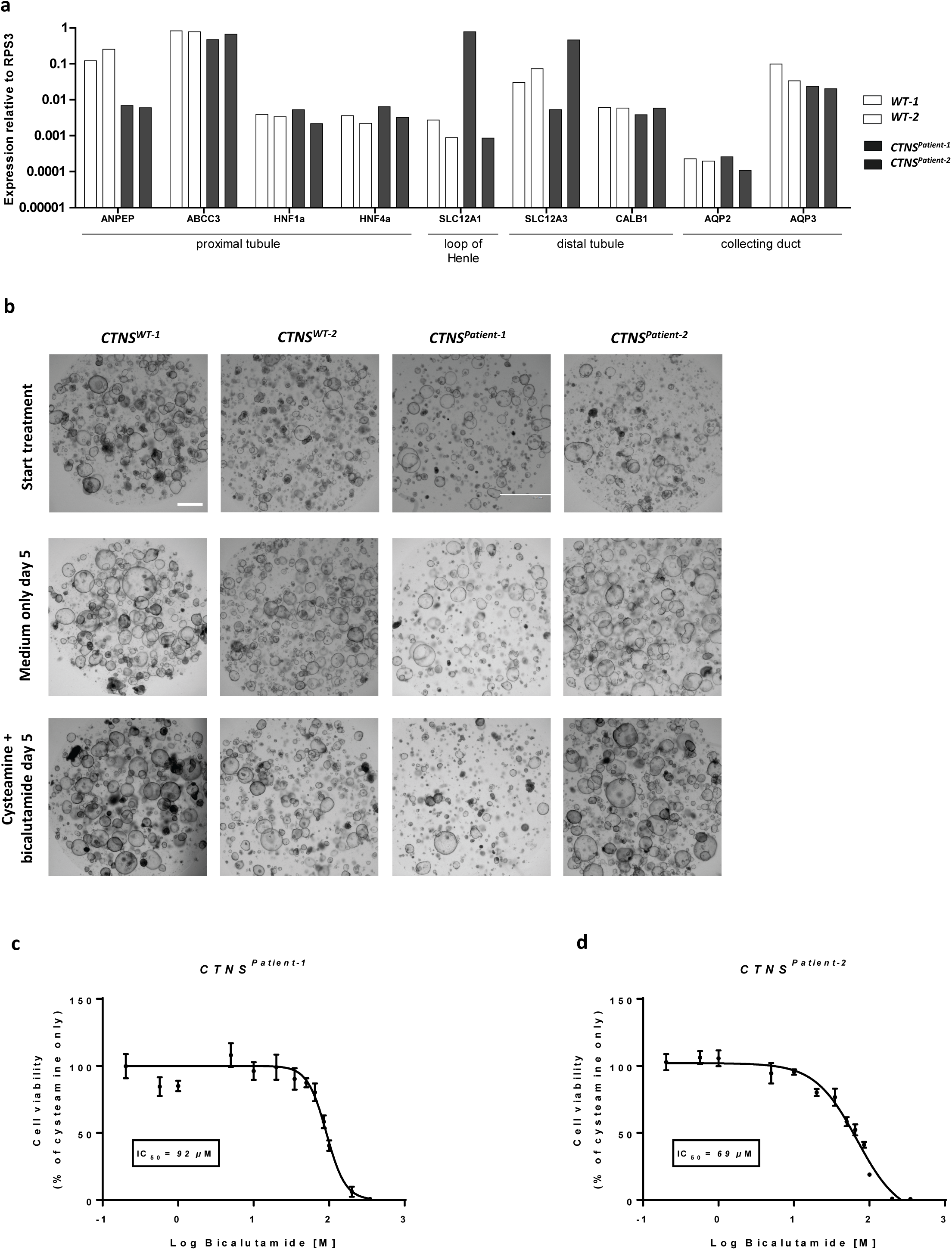
Cysteamine-bicalutamide combination treatment is safe in patient-derived cystinotic kidney tubuloids. **a**, Patient-derived cystinotic tubuloids (*CTNS^Patient-1^* and *CTNS^Patient-2^*) and tubuloids established from healthy kidney tissue (*CTNS^WT-1^* and *CTNS^WT-2^*) were differentiated for 7 days and analyzed by quantitative real-time PCR for markers of various segments of the nephron. **b**, Brightfield images of cystinotic tubuloids and healthy control tubuloids at the start of treatment and after 5 days of cysteamine (100 μM)-bicalutamide (35 μM) combination treatment or treatment with medium only (negative control). Scale bars are 2000 µm. **c**, **d**, Bicalutamide safety screening in control and cystinotic tubuloids. Tubuloid viability upon treatment with cysteamine (100 μM) in combination with increasing concentrations of bicalutamide was compared to treatment with cysteamine alone (= 100% viability). Data are expressed as the mean ± SEM of four replicates within one experiment. Statistical analysis was performed using one way analysis of variance (ANOVA) followed by Dunnett’s multiple comparisons test. Where appropriate unpaired two-tailed Student’s t-test were used. P-values < 0.05 were considered to be significant.

**Supplementary figure S6.**
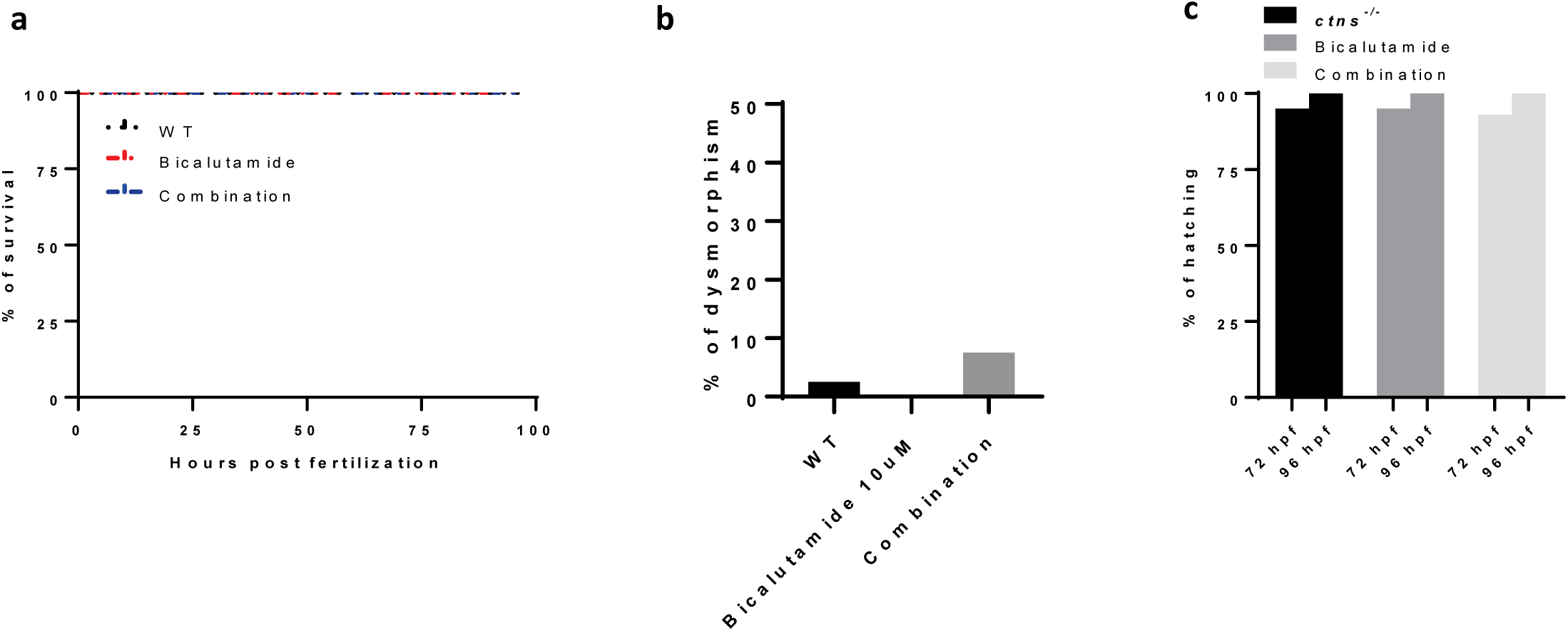
Cysteamine-bicalutamide combination treatment is safe in wild type zebrafish. **a**, Survival rates in wild type zebrafish upon treatment with bicalutamide (10 µM), and a combination of cysteamine and bicalutamide (1000 µM and 10 µM, respectively). **b**, Deformity rates in wild type zebrafish upon treatment with bicalutamide (10 µM), and a combination of cysteamine and bicalutamide (1000 µM and 10 µM, respectively). **c**, Hatching rates in surviving wild type zebrafish evaluated at 72 and 96 hrs post fertilization (hpf) with bicalutamide (10 µM), and a combination of cysteamine and bicalutamide (1000 µM and 10 µM, respectively). The total numbers of embryos evaluated for survival, hatching, and deformity rates were 40 embryos per group. Drugs were administered at 48 hpf in all experiments dissolved in the swimming water with the specified concentrations. Data are expressed as the mean ± SEM of three independent experiments. Statistical analysis was performed using one way analysis of variance (ANOVA) followed by Dunnett’s multiple comparisons test. Where appropriate unpaired two-tailed Student’s t-test were used. P-values < 0.05 were considered to be significant.

**Supplementary file 1 and 2.** Metabolomics data of ciPTEC and tubuloids, respectively.

**Supplementary Table 1.**
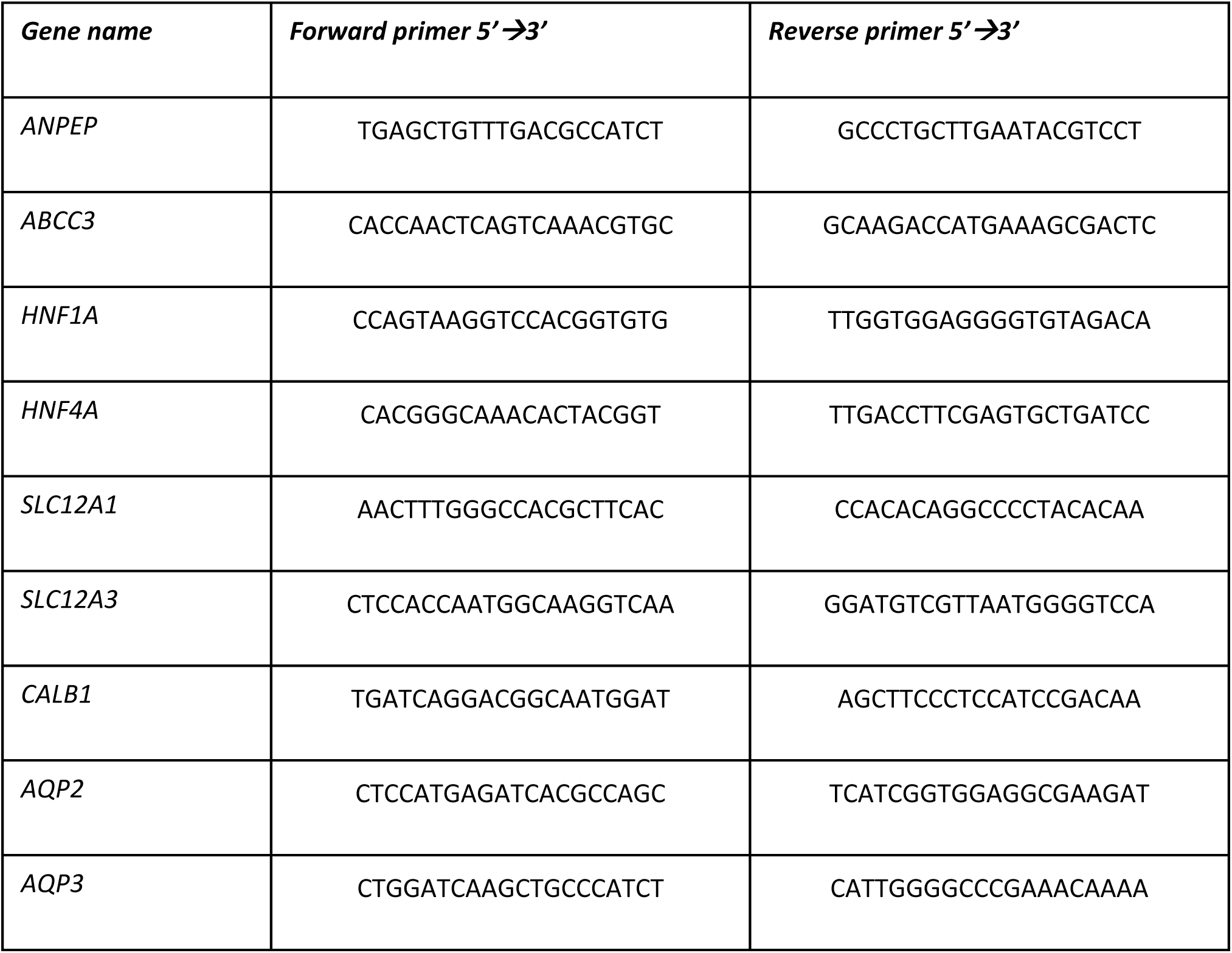
Sequences of the primers used for tubuloids study.

